# Population genomic and evolutionary modelling analyses reveal a single major QTL for ivermectin drug resistance in the pathogenic nematode, *Haemonchus contortus*

**DOI:** 10.1101/298901

**Authors:** Stephen R. Doyle, Christopher J. R. Illingworth, Roz Laing, David J. Bartley, Elizabeth Redman, Axel Martinelli, Nancy Holroyd, Alison A. Morrison, Andrew Rezansoff, Alan Tracey, Eileen Devaney, Matthew Berriman, Neil Sargison, James A. Cotton, John S. Gilleard

**Affiliations:** Wellcome Sanger Institute, Hinxton, Cambridgeshire, CB10 1SA, United Kingdom; Stephen R. Doyle:; Alan Tracey:; James A. Cotton:; Nancy Holroyd:; Matthew Berriman:; Axel Martinelli; Department of Genetics, University of Cambridge, Downing Street, Cambridge CB2 3EH, United Kingdom; Chris J. R. Illingworth: (also Department of Applied Maths and Theoretical Physics, Wilberforce Road, Cambridge CB3 OWA); Institute of Biodiversity Animal Health and Comparative Medicine, College of Medical, Veterinary and Life Sciences, University of Glasgow, Garscube Campus, Glasgow, G611QH, United Kingdom; Roz Laing:; Eileen Devaney; Moredun Research Institute, Pentlands Science Park, Bush Loan, Penicuik EH26 OPZ, United Kingdom; David J. Bartley:; Alison A. Morrison; Department of Comparative Biology and Experimental Medicine, Faculty of Veterinary Medicine, University of Calgary, Calgary, Alberta, Canada; John S. Gilleard:; Elizabeth Redman:; Andrew Rezansoff; University of Edinburgh, Royal (Dick) School of Veterinary Studies, Edinburgh, EH25 9RG, United Kingdom; Neil Sargison

**Keywords:** *Haemonchus contortus*, ivermectin, drug resistance, genome sequencing, population genetics, genetic mapping

## Abstract

**Background:** Infections with helminths cause an enormous disease burden in billions of animals and plants worldwide. Large scale use of anthelmintics has driven the evolution of resistance in a number of species that infect livestock and companion animals, and there are growing concerns regarding the reduced efficacy in some human-infective helminths. Understanding the mechanisms by which resistance evolves is the focus of increasing interest; robust genetic analysis of helminths is challenging, and although many candidate genes have been proposed, the genetic basis of resistance remains poorly resolved.

**Results:** Here, we present a genome-wide analysis of two genetic crosses between ivermectin resistant and sensitive isolates of the parasitic nematode *Haemonchus contortus*, an economically important gastrointestinal parasite of small ruminants and a model for anthelmintic research. Whole genome sequencing of parental populations, and key stages throughout the crosses, identified extensive genomic diversity that differentiates populations, but after backcrossing and selection, a single genomic quantitative trait locus (QTL) localised on chromosome V was revealed to be associated with ivermectin resistance. This QTL was common between the two geographically and genetically divergent resistant populations and did not include any leading candidate genes, suggestive of a previously uncharacterised mechanism and/or driver of resistance. Despite limited resolution due to low recombination in this region, population genetic analyses and novel evolutionary models supported strong selection at this Q.TL, driven by at least partial dominance of the resistant allele, and that large resistance-associated haplotype blocks were enriched in response to selection.

**Conclusions:** We have described the genetic architecture and mode of ivermectin selection, revealing a major genomic locus associated with ivermectin resistance, the most conclusive evidence to date in any parasitic nematode. This study highlights a novel genome-wide approach to the analysis of a genetic cross in non-model organisms with extreme genetic diversity, and the importance of a high quality reference genome in interpreting the signals of selection so identified.

## Background

Parasitic worms, commonly termed helminths, are extremely diverse and frequently responsible for significant morbidity and mortality in their hosts. Control of helminths of human and veterinary importance is heavily dependent on the large-scale administration of anthelmintic drugs; for instance, the macrocyclic lactone ivermectin has been extremely successful in the control of a number of helminths in both humans and animals [1]. These successes are now threatened by the emergence of drug resistance. In many species of parasitic nematodes of livestock, anthelmintic drug resistance is already a major problem for global agricultural production and animal welfare and the rapid acquisition of resistance to both single and multiple drug classes has been widely documented [2, 3]. Furthermore, there are growing concerns regarding the reduced efficacy of compounds used in mass drug administration (MDA) programs for some human-infective helminths, which may reflect the emergence of resistance [4–7]. In spite of the importance of this issue, remarkably little is known regarding the molecular mechanisms of resistance to most anthelmintic drug groups, with the notable exception of the benzimidazole class. This is, at least in part, due to a lack of genomic resources, tools and techniques with which to study these experimentally challenging organisms.

*Haemonchus contortus* is a gastrointestinal parasite of wild and domesticated ruminants that has a major impact on the health and economic productivity of sheep and goats globally. Resistance of *H. contortus* to almost all of the classes of anthelmintic drugs, including to multiple classes simultaneously, has been documented in many regions of the world [8–12], and can arise within a just a few years of introduction of a new drug class [13, 14]. Partly for these reasons, *H. contortus* has emerged as a model parasitic nematode to characterise anthelmintic resistance, as well as drug and vaccine discovery research as alternate means of control [15–17]. Its utility as a model is largely due to a greater amenability to experimentation than most parasitic nematodes; it is possible to establish and maintain isolates *in vivo* in the natural host, perform genetic crosses *in vivo*, and undertake *in vitro* culture for part of its life cycle, allowing drug assays and genetic manipulation such as RNAi to be performed [18]. The ability to utilise these molecular approaches is complemented by extensive information about the structure of the genome and transcriptional differences between the major life stages [19–21].

Research into the genetic basis of ivermectin resistance has been dominated by the examination of a number of candidate genes [22]. Chosen based on their potential roles in the mechanism of action or efflux of drugs, many candidate genes have been proposed to be associated with ivermectin resistance in *H. contortus* (and other parasitic nematodes targeted with ivermectin) on the basis of studies comparing SNP or haplotype frequencies between small numbers of resistant and susceptible isolates [23–27], or pre- and post-ivermectin treatment [28, 29]. However, considering the extremely high levels of genetic diversity of *H. contortus* populations, together with the limited number of well-characterised ivermectin resistant isolates, at best only circumstantial and inconsistent support is available for the involvement of any of the leading candidate genes. Further, careful validation of a number of these candidates in different field populations [30] or in controlled crosses [31] has failed to support previously defined associations between tested candidates and their associated resistance profile. The lack of consistency between studies had led to much discussion and debate regarding the complexity of the genetic basis of ivermectin resistance, both in terms of the number of loci involved and the extent of genetic variation between geographically distinct resistant isolates [22, 32, 33]. Moreover, the discordance among studies broadly reflects the difficulty in identifying real associations between genotype and phenotype simply by comparing resistant and susceptible populations, or individuals, due to the confounding effects of extremely high levels of genome-wide genetic variation [34].

Genome-wide and genetic mapping approaches are emerging for *H. contortus* due to progress in genetic crossing methodologies [21, 35–40] and an increasing complement of genomic resources [19–21]. Two serial backcrosses have previously been undertaken between the susceptible isolate MHco3(ISE) and two geographically independent ivermectin resistant isolates, MHco4(WRS) and MHco10(CAVR) [39] (**Fig 1 A,B**). Co-segregation of a single microsatellite marker, Hcms8a20, confirmed that these backcrosses had successfully introgressed ivermectin resistance loci from the two resistant populations into the susceptible MHco3(ISE) genomic background. Subsequent genetic studies showed that none of the leading candidate genes from the literature - *Hco-avr-14, Hco-glc-5, Hco-lgc-37, Hco-pgp-9, Hco-pgp-2* and *Hco-dyf-7 -* showed evidence of introgression [31], and therefore, the genetic mediator(s) of resistance remain unresolved. However, the increasing accessibility to high throughput sequencing of non-model organisms, together with a high quality reference genome for *H. contortus*, offers an opportunity to characterise precisely the genome-wide evidence of introgression and genetic architecture in this parasite.

**Fig. 1.**
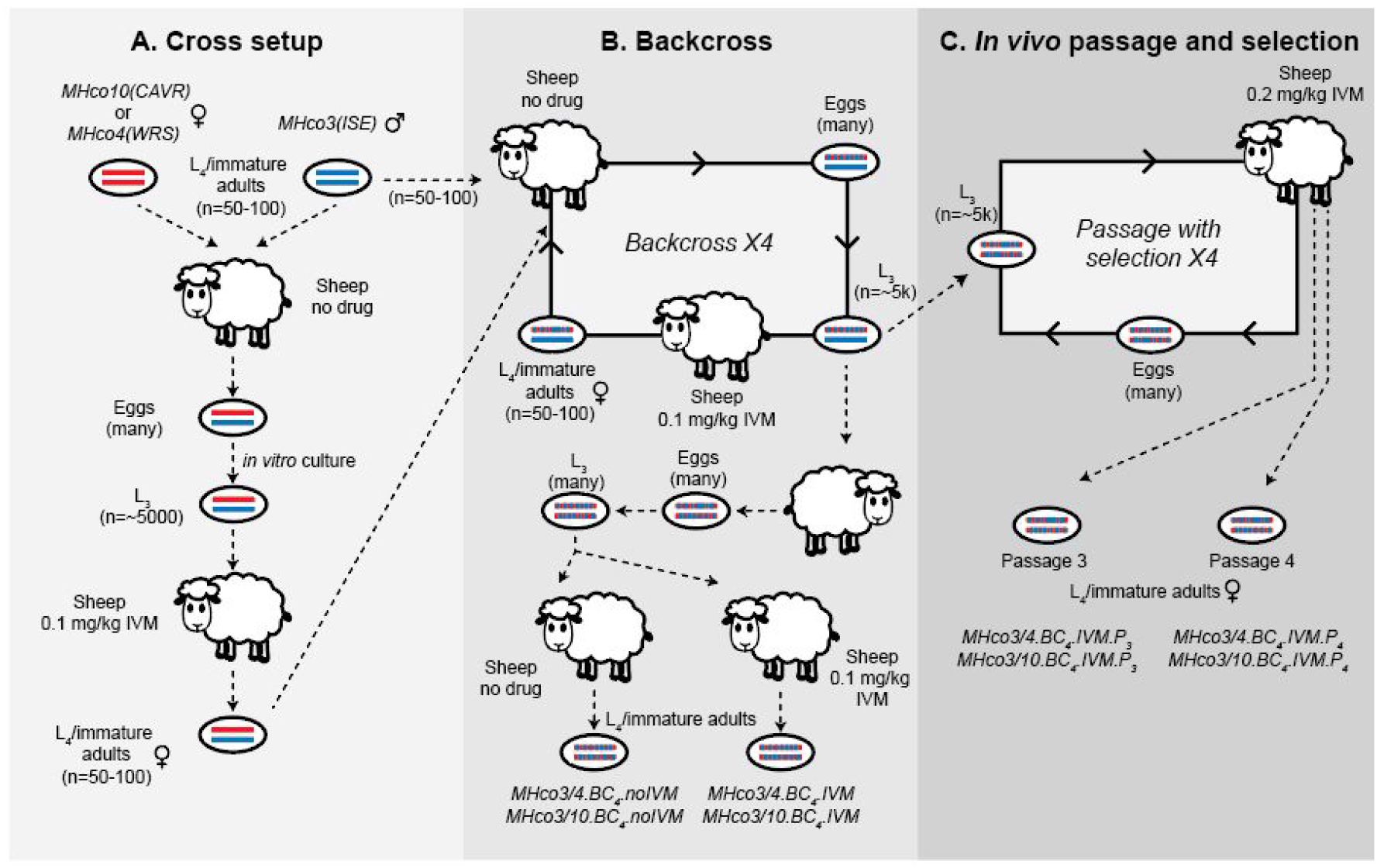
Outline of the initial crosses, backcrosses, *in vivo* passage and selection experiments. **A.** Ivermectin resistant populations MHco10(CAVR) or MHco4(WRS) (“resistant” haplotypes are depicted as red lines) were crossed with an ivermectin sensitive population MHco3(ISE) (“susceptible” haplotypes as blue lines) to generate heterozygous F_1_ progeny. F_1_ eggs were cultured *in vitro* to L_3_, before maturation *in vivo* to L_4_/immature adults, from which females were used to initiate the backcross. **B.** The first round of backcross was performed by crossing heterozygous females from the initial cross with susceptible MHco3(ISE) males *in vivo*, resulting in F_2_ progeny with reduced heterozygosity due to enrichment of MHco3(ISE) haplotypes. Resistance alleles were maintained in the backcross population by selection with ivermectin, before seeding the next round of the backcross with cross-derived heterozygous females and new susceptible MHco3(ISE) males. This process was repeated for four rounds of backcrossing, resulting in the backcrossed population becoming genetically similar to the MHco3(ISE) parental line in all regions of the genome not linked to ivermectin resistance. After four rounds of backcrossing, introgressed L_3_ progeny were used to infect a new recipient sheep, resulting in segregation of susceptible and resistant alleles in both haplotypes among the progeny (mixed red/blue haplotypes). Eggs were cultured to L_3_ before infecting two sheep, one exposed to ivermectin and one that did not receive drug. Post ivermectin treatment, L_4_/immature adults from both sheep were recovered on necropsy for sequencing. C. Post-backcross L_3_ were further passaged into a worm-naive sheep and treated with ivermectin, after which eggs were recovered and cultured to infective L_3_ for reinfection. This process was repeated for four generations of passage with selection (but without backcrossing). L_4_/immature adults were recovered after passage three and four for sequence analysis.

Here we build upon these two previous genetic crosses, extending them for further generations of passage with ivermectin selection (**Fig 1 C**), and generating whole genome sequencing data throughout the experiment. We used a bulk segregant approach, together with a recently improved chromosome-scale genome assembly, to characterise the genetic architecture in the two genetically and geographically distinct ivermectin resistance populations of *H. contortus*. In a bulk segregant analysis, allele frequency differences are determined genome-wide between pools of individuals that differ in a defined phenotype. This approach has recently been used to map resistance-associated genes in a number of field and laboratory crosses of parasites [41–43], including to map variation associated with ivermectin efficacy in three different helminth species [5, 44, 45]. We analyse these data with traditional population genetic and novel statistical methods to identify and characterise a single genomic introgression region and major QTL associated with resistance. This work represents the most comprehensive analysis of the genome-wide impact of selection and genetics of anthelmintic resistance in a parasitic nematode to date. Further, the study demonstrates the power of using a genetic crossing approach, enhanced by the use of a highly contiguous genome assembly as a framework for genome-wide analyses, to eliminate false positive genetic signals in candidate genes previously associated with resistance.

## Results

### Extensive genetic diversity defines the parental *Haemonchus contortus* populations

Whole genome sequencing of two ivermectin resistant isolates (MHco10[CAVR] and MHco4[WRS]) and one susceptible isolate (MHco3[ISE]) (**Fig 2B**) revealed high levels of nucleotide diversity throughout the five autosomes of the genome (**Fig 2A**), with the average diversity almost twice as high in the resistant isolates (MHco10[CAVR] mean *π* = 0.035 ± 0.008 standard deviations (SD); MHco4[WRS] mean *π* = 0.038 ± 0.008 SD) than the susceptible one (MHco3[ISE] mean *π* = 0.022 ± 0.008 SD) (**Fig 2C; S1 Fig A**). The two X chromosomal scaffolds were significantly less diverse (MHco3[ISE] mean *π* = 0.008 ± 0.008, MHco10[CAVR] mean *π* = 0.014 ± 0.012, MHco4[WRS] mean *π* = 0.017 ± 0.014) (**Fig 2C; S1 Fig A**), consistent with our recent finding that the X chromosome contains as little as 10% as much genetic diversity relative to the autosomes [21]. Each parental population contained local regions of high diversity that differentiated it from the others; of the ~7.6 million biallelic SNPs distributed in the genomes of the samples analysed, ~514 thousand were private to MHco3(ISE) (**S2 Fig A**), ~960 thousand to MHco10(CAVR) (**S2 Fig B**) and ~685 thousand to MHco4(WRS) (**S2 Fig C**). In addition, high quality homozygous structural variants were evident between populations (**S3 Fig A,C,E**), with a large number of deletions, and substantially fewer duplications and inversions detected (**S3 Fig B,D,F**). Further, we examined short-range haplotype diversity in each population by measuring linkage disequilibrium (LD) between pairs of SNPs detected in paired reads (**S1 Fig B**). Although restricted by the Pool-seq design to pairwise SNP comparisons less than 500 bp apart, i.e., within a read pair, we nevertheless observed considerable decay in LD over this distance. Moreover, the rate of LD decay was correlated with nucleotide diversity: a greater loss of LD in the MHco4(WRS) and MHco10(CAVR) populations over the 500 bp was observed, relative to the less diverse, susceptible MHco3(ISE) population.

**Fig. 2.**
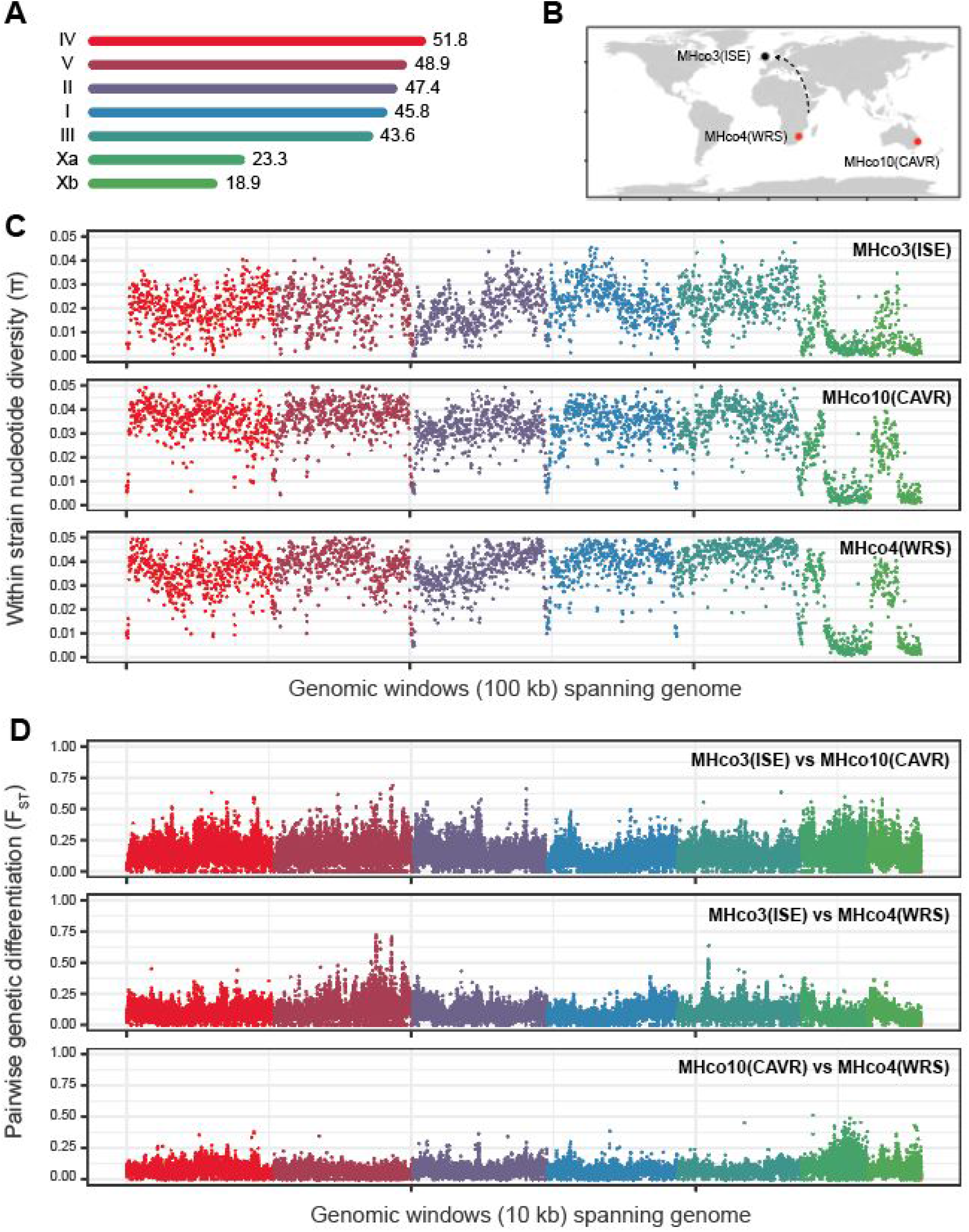
Genetic diversity within and between parental populations used in the genetic cross. **A.** The 279 Mbp V3 genome assembly of *H. contortus* consists of five autosomal and two X chromosome scaffolds, named based on synteny with *Caenorhabditis elegans* chromosomes. The size of each scaffold is indicated, and are presented in order by length (Mbp). **B.** Geographic origin of the susceptible MHco3(ISE) and ivermectin resistant MHco1O(CAVR) and MHco4(WRS) populations used in the genetic crosses. All populations are archived at the Moredun Research Institute, UK – while MHco3(ISE) has been maintained there for decades, it was originally isolated in East Africa. **C.** Within-population nucleotide diversity for each of the parental populations, calculated as mean diversity per 100 kbp windows throughout the genome using *npstat*. **D.** Between population diversity, calculated as pairwise F_ST_ in 10 kbp windows throughout the genome using *popoolation2*. Colours here and throughout represent the chromosomal scaffolds as described in A.

Genetic diversity between each of the three parental populations was assessed by pairwise analysis (measured by F_ST_ of single nucleotide polymorphisms [SNPs] in 10 kbp windows), which confirmed significant genome-wide differentiation (**Fig 2D**). This is highlighted by multiple discrete peaks of differentiation within each chromosome, both between resistant and susceptible stains, but also between the two resistant populations. We must therefore conclude that most of this genetic differentiation reflects underlying genetic structure between populations unrelated to their drug resistance phenotype; this is perhaps not surprising, given they are derived from reproductively isolated populations due to their geographic distribution (**Fig 2B**). In addition, MHco3(ISE) was subject to multiple rounds of inbreeding during is original derivation as a laboratory population [46], whereas the resistant populations have not been subjected to deliberate inbreeding and were isolated from outbred populations more recently. Collectively, these data emphasise the challenge of characterising genetic variation associated with phenotypes such as drug resistance by simply comparing genetic diversity between a susceptible and resistant population without accounting for the extensive background genetic variation present.

### Genome-wide analysis of genetic diversity reveals the same single large introgressed region in both backcross lines

To map genetic variation linked to ivermectin resistance in the two parental resistance populations, we sequenced pre- and post ivermectin treatment samples of the initial backcross between each of MHco10(CAVR) and MHco4(WRS) populations with the MHco3(ISE) susceptible genetic background [39], as well as after subsequent passage of the introgressed populations with further treatment. Consistent with the backcross design, pairwise comparisons were made between specific stages of the experiment and the MHco3(ISE) susceptible parent. We hypothesised that where backcross populations were not subjected to further ivermectin treatment (**Fig 1**: MHco3/10.BC_4_.noIVM & MHco3/4.BC_4_.noIVM), there would be relatively little genetic differentiation throughout the majority of the genome between the defined population and MHco3(ISE). By contrast, where populations were subjected to further ivermectin treatment, we hypothesised that genetic differentiation with MHco3(ISE) should increase close to any resistant allele, with high levels of differentiation in regions of the genome linked to an ivermectin resistance-conferring locus and low levels of differentiation elsewhere.

As hypothesised, little genetic differentiation between the post-backcross, no-selection population and the susceptible parental population was observed in either cross (**Fig 3**; MHco3/10.BC_4_.noIVM and MHco3/4.BC_4_.noIVM). After backcrossing, in the absence of selection, genetic material from the susceptible parent should comprise approximately 97% of the population due to the repeated backcrossing with MHco3(ISE) (resistant alleles comprise half of the initial cross population and reduce by a half with each backcross), making them largely indistinguishable from the susceptible parent. Selection increases the fraction of alleles derived from the resistant parent, but is limited by the backcross design; no more than 50% of the genetic material at any location may originate from the resistant parent. Upon further selection (**Fig 3**; MHco3/10.BC_4_.IVM and MHco3/4.BC_4_.IVM), a region of differentiation from the susceptible parent, particularly in the MHco3/10 cross, was localised to the right arm of chromosome V. This region was markedly differentiated in both MHco3/10 and MHco3/4 crosses after *in vivo* selection and passage (**Fig 3**; MHco3/10.BC_4_.IVM.P_3_, MHco3/4.BC_4_.IVM.P_3_, MHc3/10.BC_4_.IVM.P_4_ and MHc3/10.BC_4_.IVM.P_4_). No other region of the genome displayed a marked and progressive increase in differentiation throughout the crosses. To test this explicitly, we determined the correlation between change in F_ST_ per each genomic window and progression through the cross; only the chromosome V region demonstrated progressive genetic differentiation over time (**S4 Fig A,B**), which was highly correlated between the two crosses (**S4 Fig C**).

**Fig. 3.**
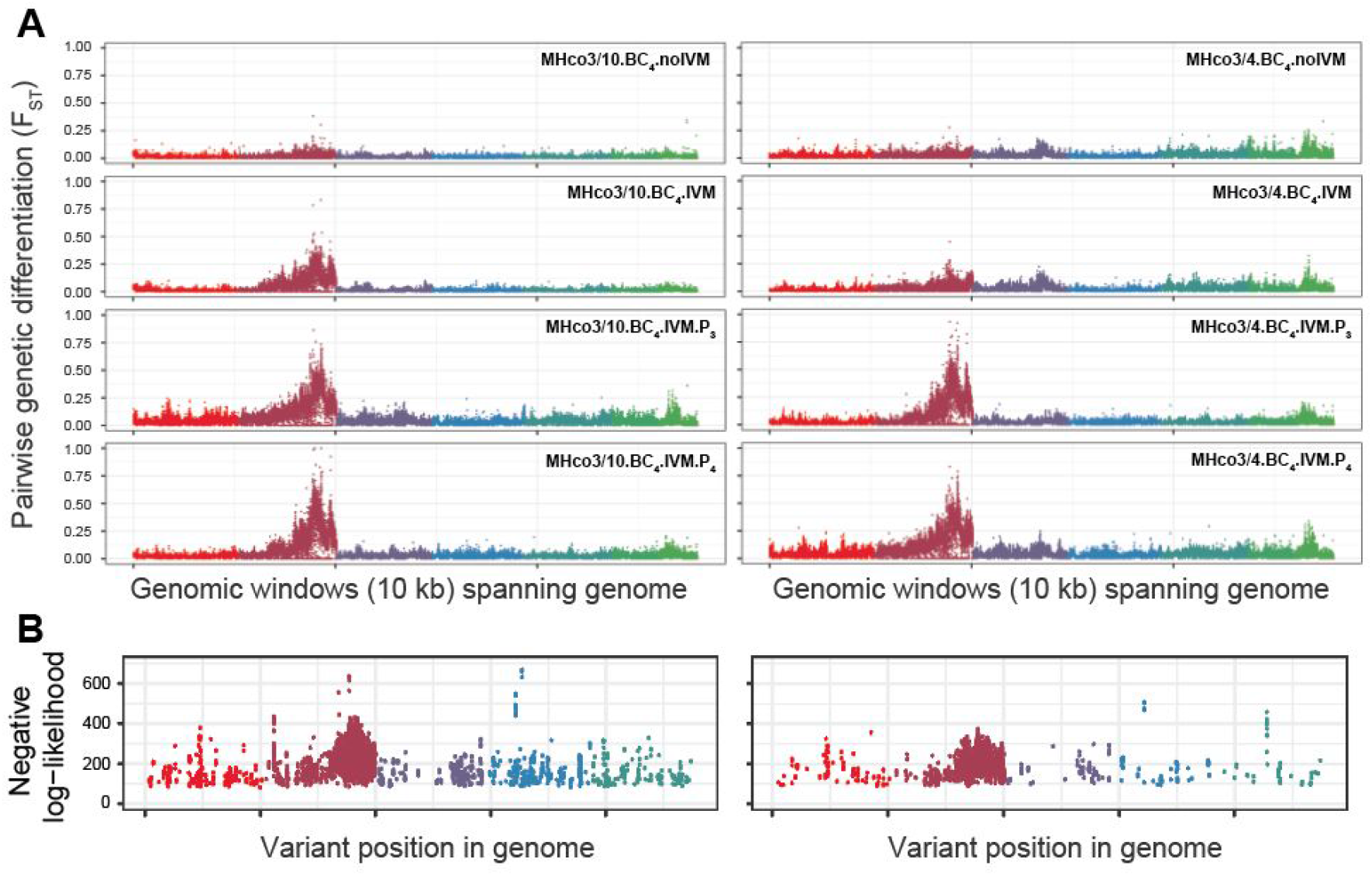
Genome-wide analysis of ivermectin associated loci. **A.** Pairwise genetic diversity throughout the MHco3/10 (left plots) and MHco3/4 (right plots) backcrosses. In both crosses and at each point in the backcross, the experimental population was backcrossed against the MHco3(ISE) parental population – the data are therefore presented to compare the genetic diversity (F_ST_ measure in 10 kbp windows using *popoolation2*) between MHco3(ISE) and each sampled time point in the cross as per the cross scheme in Fig 1. **B.** Output of the single-locus evolutionary model, describing sites which were inconsistent with a model of neutral evolution, measured using a likelihood threshold. Note that the X chromosome is not represented. Colours represent the chromosomal scaffolds as described in Fig 2A.

Collectively, these data suggest that there is a single introgression region on chromosome V representing a major QTL that is under selection upon drug exposure, and that this region is under selection in the two independent serial backcrosses performed. This conclusion was supported by an independent evolutionary analysis, which compared the observed frequencies of variants associated with the resistant parental population with the distribution of allele frequencies expected under the assumption of selective neutrality (**S6 Fig**). An over-representation of alleles (71% and 91% of the total number of such alleles for the MHco3/10 (**S6 Fig A**) and MHco3/4 (**S6 Fig B**) crosses, respectively) with atypical frequencies towards the right hand end of chromosome V were strongly inconsistent with the neutral expectation, suggesting that they had increased in frequency under the influence of linked selection (**Fig 3B**).

### Characterisation of the introgression region located on chromosome V

A comparison of genetic diversity on chromosome V between MHco3(ISE) and the end-point of both crosses (BC_4_.IVM.P_4_) revealed a strikingly similar pattern between the crosses, with a major region of differentiation spanning 37-42 Mbp, and a second but lesser increase in genetic differentiation between 45-48 Mbp that is most prominent in the MHco3/10 population (**Fig 4A**). Encouragingly, the Hcms8a20 microsatellite marker (located at position 36.16 Mbp along chromosome V) that was shown to be genetically linked to ivermectin resistance in the preliminary analysis of these backcrosses [39] lies adjacent to the QTL region, suggesting that while it is strongly linked to resistance, it is unlikely to be completely linked with a locus directly causing this phenotype. The F_ST_ data are supported by a comparison of Tajima’s D throughout this region (**Fig 4B**; top plots). Although variation in Tajima’s D was observed throughout the genome in both crosses (**S7 Fig**), the greatest variation between the susceptible parent and backcrossed lines was observed at approximately 39-40 Mbp (**Fig 4B**; bottom plots), which was associated with significantly negative Tajima’s D in the backcrossed lines consistent with strong positive selection. In contrast to the F_ST_ data, analysis using Tajima’s D suggested a single peak, corresponding to a single site or region under positive selection.

**Fig. 4.**
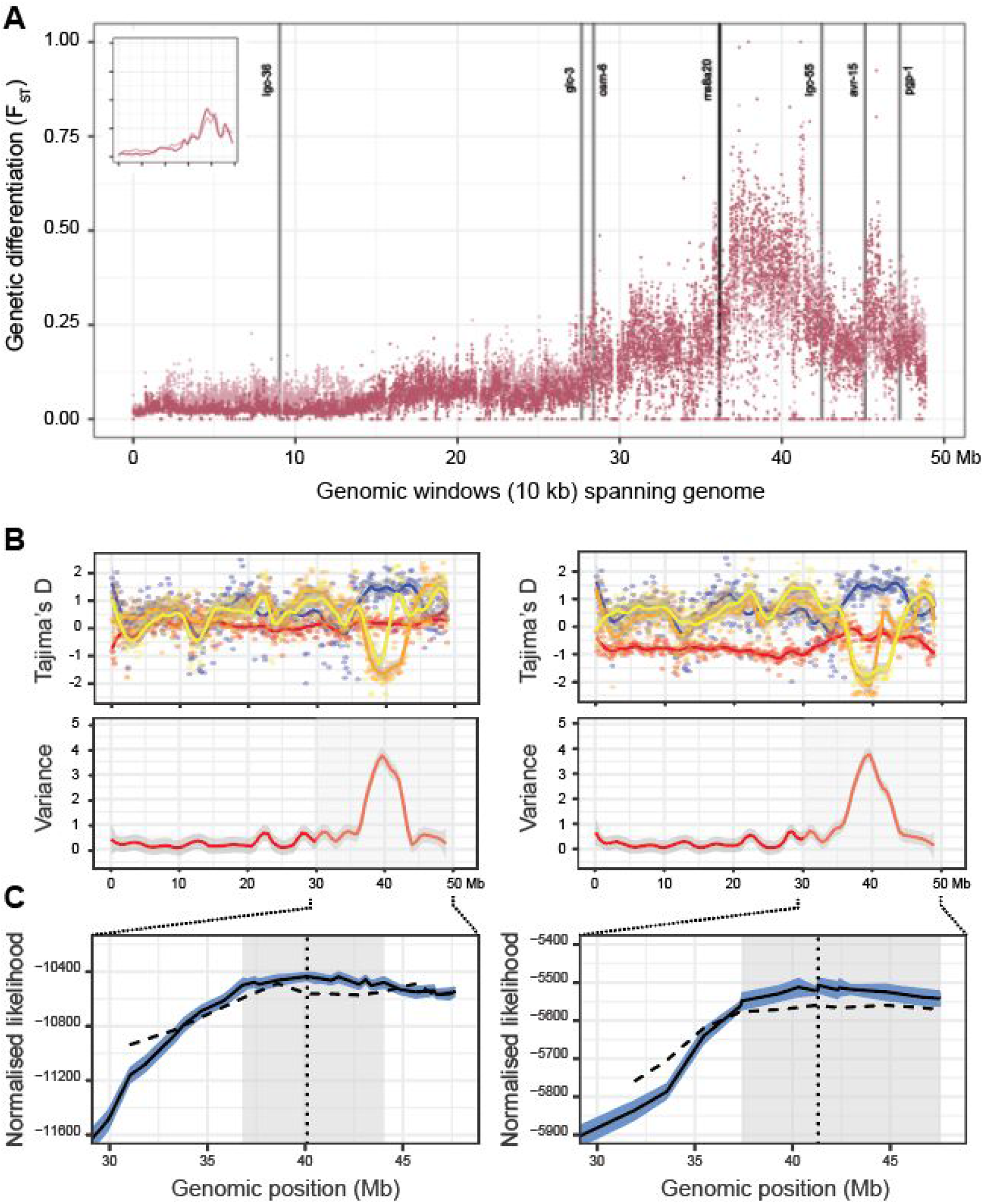
Analysis of chromosome V and the major region of introgression. **A.** Genetic differentiation (F_ST_) measured in 10 kbp windows throughout chromosome V is presented between MHco3(ISE) parent and both MHco3/10.BC_4_.IVM.P_4_ (light) and MHco3/4.BC_4_.IVM.P_4_ (dark) representing the final time point of the crosses. Inset presents the smoothed F_ST_ distribution of the two comparisons. Published candidate genes in or near the introgression region (grey vertical lines), as well as the original Hcms8a20 microsatellite marker that Redman *et al*. (2012) linked to ivermectin resistance (black vertical line), are presented. **B.** Comparison of Tajima’s D per chromosome between MHco3(ISE) parent (blue), MHco10(CAVR) (left panels) or MHco4(WRS) (right panels) and passage 3 (MHco3/10.BC_4_.IVM.P_3_ or MHco3/4.BC_4_.IVM.P_3_; orange) and passage 4 (MHco3/10.BC_4_.IVM.P_4_ or MHco3/4.BC_4_.IVM.P_4_; yellow) of the crosses (top panels). Tajima’s D was calculated using *npstat* in 100 kbp windows spanning the genome. To emphasise the difference between the susceptible MHco3(ISE) and resistant passages 3 and 4 samples, we calculated the variance in the mean value of Tajima’s D between samples, which is presented as red smoothed line (bottom panel). The null expectation is that variance between these samples will be low in regions of the genome not under selection. **C.** Inferred likelihoods from the multi-locus population genetic model. The mean likelihood of the best fitting single-locus model at each locus position is shown by the solid black line; the light blue interval around this line shows the 5-95% confidence interval for this statistic calculated by a bootstrapping process. The position of the maximum likelihood value is shown by the vertical black dotted line; a confidence interval for this position, calculated from the bootstrapping values, is shaded in gray. The mean likelihood of the best fitting constrained two-locus model is shown by the black dashed line; the value shown represents the best value of the model given that one of the selected loci is at the given position.

A total of 74,424 variants (4.71% of the total variants found on chromosome V) were found to be segregating within the 37-42 Mbp region, of which 14,009 and 12,827 variants presented a signal of differentiation (greater than five standard deviations from the genome-wide mean P value from the Fisher’s exact test) between the susceptible and CAVR and WRS lines, respectively, due to variable degrees of linked selection with the causative allele. As the genome assembly used in this study was not annotated, it was difficult to prioritise this variant list further in the context of their impact on coding sequences. However, many candidate genes have been proposed in the literature as being in association with ivermectin resistance; to determine their possible role in driving resistance here, we curated a list of genes that have been explored in *H. contortus*, and/or had been shown to confer ivermectin resistance when mutated in *C. elegans*. We determined the location of these genes in the current assembly, either via mapping the published gene models from the V1 annotation, or determining the closest *H. contortus* orthologous gene from *C. elegans* candidates using Wormbase Parasite [47] (**S2 Table; S9 Fig**). At least one candidate gene was located in each chromosome; in chromosome V, the location of six candidate genes were determined. However, none of these genes were found in the main introgression region defined by the F_ST_ analysis (**Fig 4 A**; vertical annotated lines). Three genes – *Hco-lgc-55, Hco-avr-15*, and *Hco-pgp-1(9)*, the latter two of which have a strong association with ivermectin resistance – lie to the telomere-side of the introgression region, however, they are located on the periphery of the F_ST_ peaks rather than within them, suggesting that they are unlikely candidates to be under direct selection. Considering the absence of candidate resistance genes in the F_ST_ peaks, we conclude that the driver(s) of ivermectin resistance in the MHco4(CAVR) and MHco4(WRS) populations are novel and have not been previously described in association with ivermectin resistance.

A multi-locus population genetic model provided clarity on the extent to which the location of the selected allele could be determined by our data. In contrast to F_ST_ and Tajima’s D, this model explicitly considered linkage between variant alleles arising from their common parental origin. Putative segregating sites in chromosome V between the resistant and susceptible parental populations were identified: A total of 474 such sites were identified in the MHco3/10 dataset, with 157 sites in the MHco3/4 dataset. Calculations were then performed upon data from these loci. The maximum likelihood of the allele under selection was in a broadly consistent location in each experiment, being inferred to exist at 40.10 and 41.31 Mbp respectively. However, there was considerable uncertainty in this location; a conservative estimate gave confidence intervals of 36.86 to 44.03 Mbp and 37.41 to 47.56 Mbp in each case (**Fig 4 C**). These broad confidence intervals were explained by the population structures inferred by the model, which explicitly considered the location of crossover recombination events between the parental haplotypes throughout the experiment. Example structures generated using the maximum likelihood parameters for each population show how selection for resistance drives the accumulation of genetic material from the resistant parent through the course of the experiment (**Fig 5**); these outputs may be contrasted with equivalent data generated under a model of selective neutrality (**S10 Fig**). Across 250 replicates, a mean of 177.1 crossover recombination events were predicted in chromosome V of the final MHco3/10 population (range 120 to 224), giving a mean length of a block of parental genome of 17.64 Mbp; in the final MHco3/4 population a mean of 186.4 recombination events were seen (range 143 to 238), giving a mean parental haplotype block length of 17.06 Mbp. Under selection, blocks of genome containing the selected allele are favoured; the large size of each block reduces the precision with which the location of the allele under selection can be identified.

**Fig. 5.**
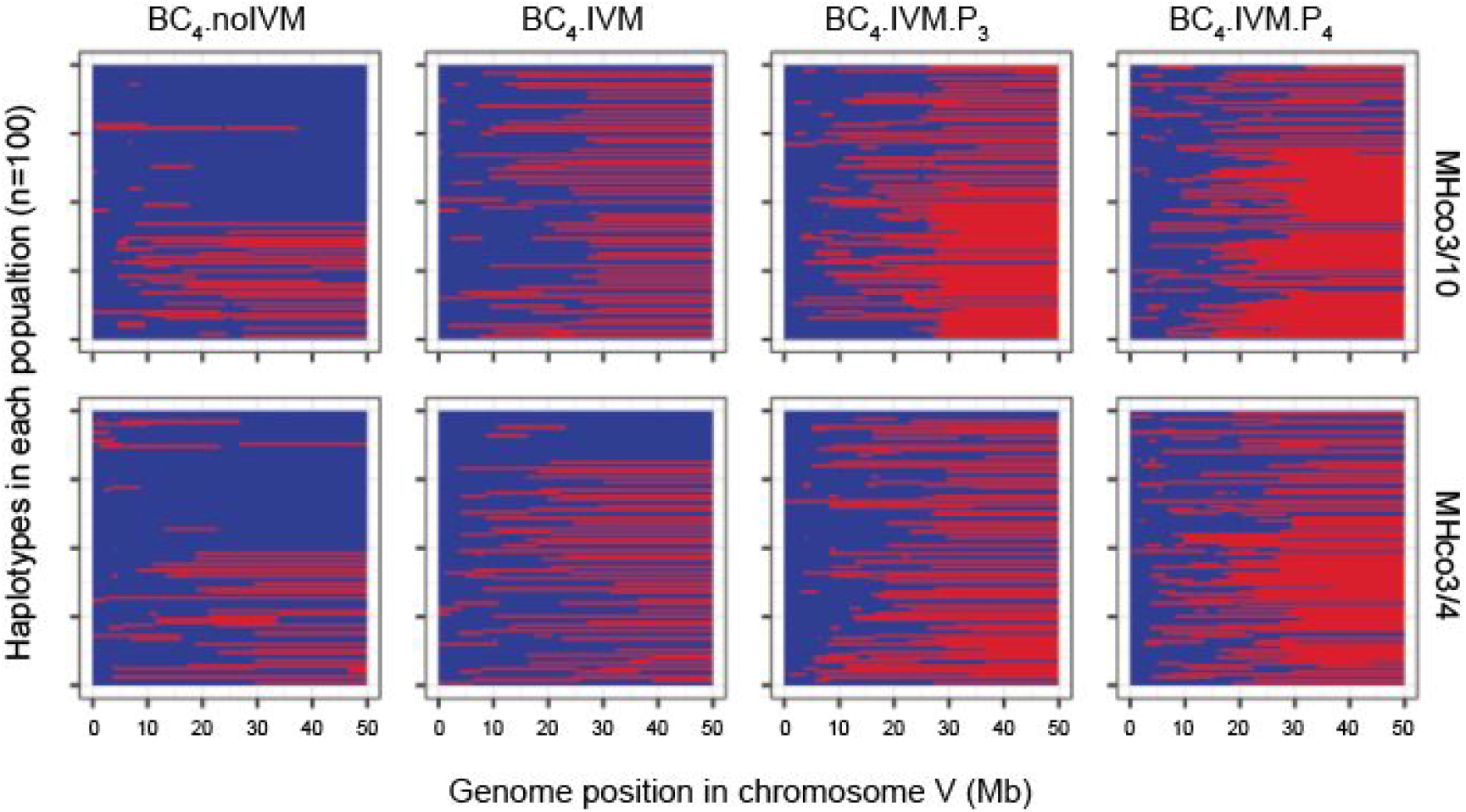
Haplotype structure of chromosome V in an example output from the multi-locus model. Segments of genome from the parent containing the resistance allele are shown in red, while segments of genome from the susceptible parent are shown in blue. Data shown were generated using the maximum likelihood parameters in each case.

An extended multi-locus model incorporating selection on alleles at two loci did not find evidence for selection at more than one allele; despite the second peak of differentiation found at approximately 45 Mbp in the F_ST_ analysis, evidence for a second site was not identified. The identification of independent selected alleles requires them to be separated in the genome by recombination; on the basis of our knowledge of recombination in this system we searched for alleles located at least 2 Mbp apart in the genome. Under this constraint, the best fitting model identified variants under selection at the positions 38.6 Mbp and 45.7 Mbp in the MHco3/10 data (**S8 Fig A**), and at the positions 41.2 Mbp and 44.8 Mbp in the MHco3/4 data (**S8 Fig B**). However, these models fit the data less well than the single-driver model described above (**Fig 4C**), favouring an explanation in which only one variant is under selection. We note that the single-driver model is as a special case of the two-driver model, for which both drivers are in precisely the same location and the selective effect of one of the drivers is set to zero; removing the constraint that the alleles be separated would result in a fit to data at least as good as the single-driver model. In conclusion, this analysis cannot exclude the presence of more than one drug-resistance variant in the identified region of chromosome V, but provides no evidence to support the existence of a second selected allele.

### Strength of selection

We further used the multi-locus single driver model to infer the strength and manner of selection in favour of the drug resistance allele. Data in each case indicated strong selection for the resistant allele (**Fig 6**), such that susceptible worms produce multiple times fewer offspring than the resistant worms under drug treatment. In each case the value of the dominance coefficient, *h*, was less than one, indicating either an additive effect for the MHco10(CAVR) case, whereby each copy of the resistance allele contributes an equal amount of resistance, or weak dominance for the MHco4(WRS) data, whereby the first resistance allele contributes slightly more than the second. This is consistent with the phenotypic characterisation of the initial crosses, whereby the F_1_ individuals that would have been heterozygous for resistant alleles were partially resistant to ivermectin [39]. As such, having two copies of the resistant allele had a greater phenotypic effect than one; these data therefore have implications for the mechanism of the drug resistance allele, and evolution of drug resistance in general terms, as discussed further below.

**Fig. 6.**
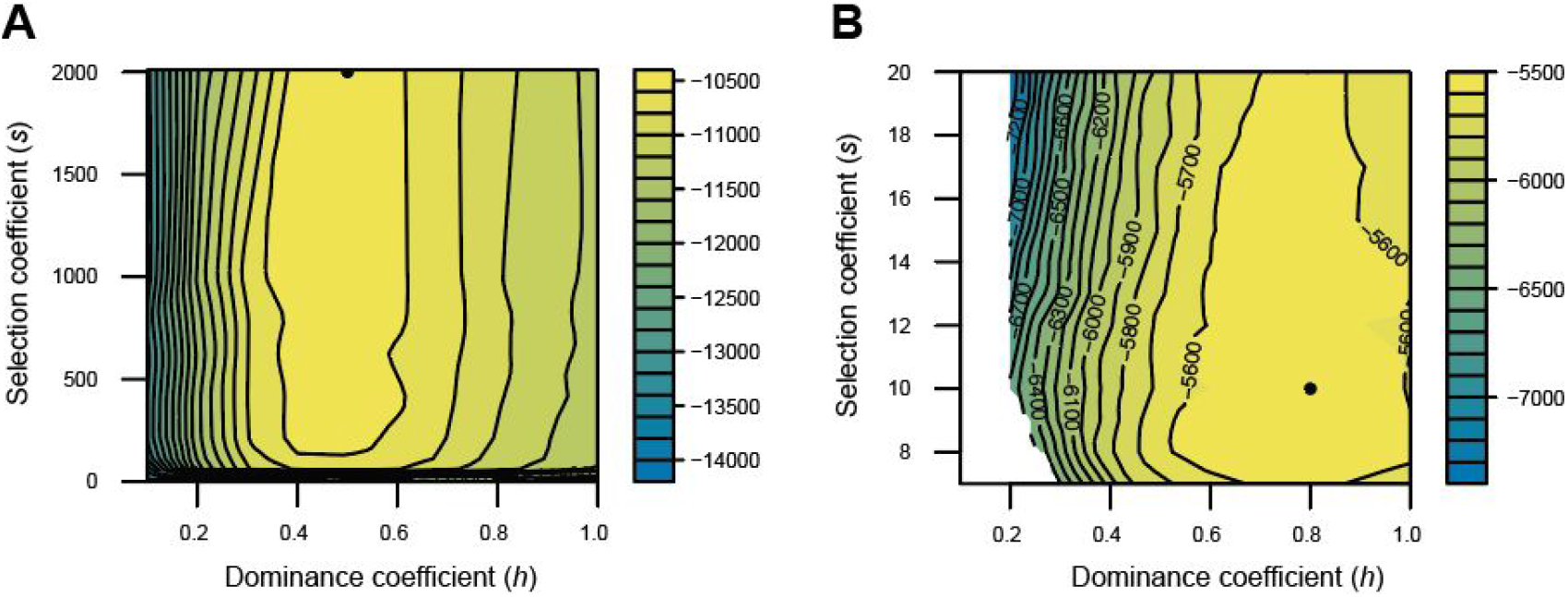
Likelihood surface describing inferred selection parameters for the resistance allele. Within our model, the fitness of a genotype is determined by the alleles at a single resistant allele, with homozygous susceptible worms having fitness 1, homozygous resistant worms having fitness 1+s, and heterozygous worms having fitness 1+*hs* in the presence of the drug. In each plot, calculated for both the parental **(A)** MHco10(CAVR) and **(B)** MHco4(WRS) populations, we identify the location of the maximum likelihood values for selection (s) and dominance coefficients (*h*) (black dot), and the likelihood landscape surrounding this maximum value (log likelihood heatmap scale: yellow = high likelihood; blue = low likelihood) for the position in the genome of the resistant allele. Changes of the order of 100 likelihood units indicate substantial differences in the extent to which the model fits to the data.

## Discussion

### Genetic mapping identifies a major ivermectin resistance QTL in two independent *H. contortus* populations from different continents

We have analysed whole genome sequence data collected from the most recent phase of a multi-generational crossing experiment conducted in populations of the parasitic worm *H. contortus*. Our analysis unequivocally identified a single region of introgression within chromosome V that contains at least one major ivermectin resistance allele. This major ivermectin resistance QTL is common to two independent ivermectin resistant populations – MHco10(CAVR) and MHco4(WRS) – that were originally isolated from Australia and Africa, respectively. This genomic region was clearly distinguished in the experimental data using different population genetic methods. Although this region in which we infer a selected allele to exist is relatively large (Conservatively ~5 Mbp from 37 to 42 Mbp, comprising 1.73% of the genome), each of our analytical approaches consistently identified the same region under selection, with no other comparable regions of introgression elsewhere in the genome in either of the two backcross experiments. The annotation of the *H. contortus* genome is ongoing, and hence we can only make a best guess estimate of the number of putative genes in this region: we predict approximately 20,000 genes in the 279 Mbp genome, which would correspond to almost 360 genes in the introgression region (~72 genes / Mbp). Considering the high diversity between parental populations, and limited recombination in this window, it is difficult to reduce this number of genes further. However, our data suggests this introgression region is the most important ivermectin resistance QTL in the MHco10(CAVR) and MHco4(WRS) populations.

There has been much discussion and debate regarding the complexity of the genetic basis of ivermectin resistance, both in terms of the number of loci involved and the extent of geographical variation [22, 32, 33]. Our results, however, strongly suggest a single locus (or potentially multiple closely linked loci) is likely to be the major effector of ivermectin resistance in these two populations. We acknowledge that we cannot discount previously described candidate genes as mediators of resistance in other populations of *H. contortus*, or in other nematode species. Moreover, our data does not exclude a driver of transcriptional regulation within the introgression region that is under selection, which itself influences expression of other genes, including candidate genes, outside of the introgression region. However, most of the leading published candidate ivermectin resistance genes including *Hco-glc-5, Hco-avr-14, Hco-lgc-37, Hco-pgp-2* and *Hco-dyf-7* were not located in or near the QTL, which is consistent with a recent study using targeted sequencing of these genes in which none showed a signal of introgression in these two backcross experiments [31]. This suggests that none of these candidate genes show evidence of selection in response to ivermectin treatment in the MHco10(CAVR) and MHco4(WRS) populations. Although we identified three other previously described candidate genes – *Hco-lgc-55, Hco-avr-15*, and *Hco-pgp-1(9)* – adjacent to the introgression region, none were found within the peak of the region suggesting that these also are unlikely to be major direct targets of ivermectin-mediated selection. This is in contrast to a recent genome-wide analysis of multi-drug resistance of a field population of *Teladorsagia circumcincta*, a gastrointestinal nematode of sheep and goats, in which a copy number variant of *Tc-pgp-9* was expanded in an ivermectin resistant isolate. Additionally, *Tc-lgc-55* was identified in a region of high differentiation between resistant and susceptible individuals [44]. Indeed, a key conclusion of this earlier work was that ivermectin resistance was likely to be highly multigenic. Given the fragmented nature of the *T. circumcincta* assembly, it is intriguing to speculate whether many of the signals observed in the earlier study, including one or both of these genes, are not direct mediators of resistance, but rather show evidence of selection due to being linked to nearby driver mutation, as is likely the case in *H. contortus* shown here.

What might account for the large number of candidate genes previously suggested to be associated with ivermectin resistance? The answer might lie in our direct comparison of genome-wide diversity between the parental populations (**Fig 2D**). The extent of genetic divergence between these populations is striking, and makes it easier to understand how particular sequence polymorphisms might be naively attributed to being associated with resistance, when in fact they simply represent genetic variation that occurs as a result of the independent evolutionary histories and lack of interbreeding between the populations. Our results highlight the challenge of interpreting simple direct comparison of genetically distinct populations, even with genomic approaches, when trying to disentangle those genetic polymorphisms underlying resistance from a complex background of genetic variation independent from the resistance phenotype of individuals from a given population. The use of controlled genetic crosses as presented here and elsewhere [31, 39, 44], whereby population genetic structure can be explicitly accounted for in the analysis, provides a powerful way to mitigate the challenge of discriminating resistance-causing alleles from background genetic variation.

### Population genetic and evolutionary modelling defined the boundaries of the ivermectin resistance QTL

Previous research has highlighted the value of novel methods in population genetics for analysing experimental cross populations [48]. In order to gain the maximum possible insight into the data collected, we extended previous statistical work [49, 50] to account for stochasticity in the experiment. Specifically, the random location of recombination events, in the context of strong selection, and genetic drift imposed by population bottlenecks in the experimental design, leads to potential variation in the final outcome of the experiment. With prior knowledge of the recombination rate [21], we used evolutionary simulations reproducing the experimental design to explore the range of experimental outcomes arising under multiple scenarios of selection; this allowed for a direct inference of evolutionary parameters.

Inference of selection from the multi-locus model suggested the presence of stronger selection for the drug resistance allele in the MHco3/10 cross than in the MHco3/4 cross (**Fig 6**). Further, although the data suggests that resistance is at least additive in both crosses, that is, heterozygotes are likely to confer some resistance and increased resistance is achieved by having two resistant alleles, there is some evidence for dominance in the MHco4(WRS) population, whereby the second resistant allele confers a lesser fitness advantage. A previous study reported resistance to be a dominant trait in the CAVR population [36] on the basis that no difference was observed in the resistance levels of heterozygote and homozygote resistance worms. Our finding of additivity arises from the evolution of the worm populations during passage following the backcross experiment (i.e., in BC_4_.IVM.P_3/4_ samples versus the BC_4_.IVM data), and particularly the fixation of alleles from the resistant parent at putative segregating sites near the inferred position of resistance. In our model, alleles can reach fixation only where there is selection in favour of the homozygous resistant type over and above that for heterozygote resistant individuals.

Our ability to resolve the precise location of the variant under selection was limited by the number of recombination events in the worm population. Precise identification of the location of an allele under selection requires that the allele not be linked to other nearby alleles in at least some fraction of individuals in the population. While the inherent biological recombination rate here has a role, population bottlenecks in each round of the cross induced by the use of only a limited number of individuals – 50-100 male and female worms per cross generation – reduced our ability to resolve the region of introgression further toward a single causative gene or variant. These bottlenecks were due to the limited number of L_4_ worms that it was possible to collect from each sheep and transplant in the successive generation of the cross. Genetic drift induced by successive population bottlenecks introduced considerable uncertainty in the outcome of the experiment, such that replicate sets of allele frequencies from our model showed considerable differences between each other (**S5 Fig**). This implies that the data from the experiment itself should be understood as the output of a stochastic process; beyond the clear large-scale patterns observable in the data and detailed above, more minor details of the output might not be seen again were the experiment to be repeated. The structure of the experiment thus imposes a limit on our ability to infer the location of a selected allele; an improved characterisation of selection would likely best be achieved by conducting further generations of cross as performed within this experimental framework to induce more recombination events, reducing the mean size of parental genomic blocks. However, such a course of action would be limited in scope were it not accompanied by the use of larger L_4_ populations, for example by the simultaneous passage of the population through multiple animals. Clearly, a larger or longer experiment would have cost and welfare implications; the statistical framework we have developed would help to design and justify such an approach.

We note the importance of linkage between sites in the analysis of data from genetic cross populations. Whereas standard metrics such as F_ST_ and Tajima’s D do not explicitly account for linkage between sites, we have here implemented methods which explicitly account for the genomic structure arising from the history of the cross population, including the stochasticity that exists in that structure due to genetic drift and random recombination. While standard metrics can identify sites of maximum differentiation in a population with great precision, neglect of the inherent stochasticity in the outcome of an experiment can lead to an overconfidence in the extent to which they provide an accurate diagnosis of the causative variant of selection.

### Importance of chromosome-level genome assemblies for genetic mapping and population genomics

The success of identifying a single region of introgression was dependent upon an improved chromosomal-scale reference genome assembly for the *H. contortus* isolate, MHco3(ISE).N1. This isolate was used in the original draft assembly of this species [19], which was derived via inbreeding of the same MHco3(ISE) population used in the backcrosses presented. Without a contiguous chromosomal-scale genome assembly, interpretation of these type of analyses can be extremely challenging; true genetic signal(s) can be obscured by technical artefacts associated with a fragmented assembly, such as short contigs lacking spatial orientation, multiple haplotypes present, collapsed paralogs, and poor resolution of repeat structures. As a consequence, both read mapping and variant calling in fragmented genomes can be suboptimal. These technical challenges are exacerbated in highly polymorphic species such as *H. contortus* and *T. circumcincta* [44], for which reference genomes were constructed from pools of individuals that, despite efforts to reduce genetic variation by inbreeding, were highly polymorphic. Perhaps the biggest problem with using a fragmented genome assembly for genome-wide analyses is the inability to determine linkage. Signals of selection will often be dispersed across multiple scaffolds of a fragmented draft genome assembly when in reality they are adjacent to each other in a single genomic region. This can potentially can give rise to the erroneous conclusion that multiple signals of selection are present, when in fact only a single selected locus exists. We illustrate these selection artefacts by presenting the sequence data we have generated here on different versions of the MHco3(ISE).N1 reference genome assembly (**Fig 7**). In the fragmented draft genome assemblies, the signals of genetic differentiation between susceptible and resistant populations were numerous and dispersed across many assembly scaffolds, suggesting that many discrete regions of the genome may be under selection (**Fig 7**; top plot). However, only after significant improvement in chromosome contiguity does the introgression region on chromosome V become evident (**Fig 7**; bottom plot). Genome contiguity (or lack of) thus will be an increasingly important factor in the interpretation of analyses exploiting unfinished draft reference genomes of non-model species, particularly as whole genome sequencing becomes cheaper and more widely adopted.

**Fig. 7.**
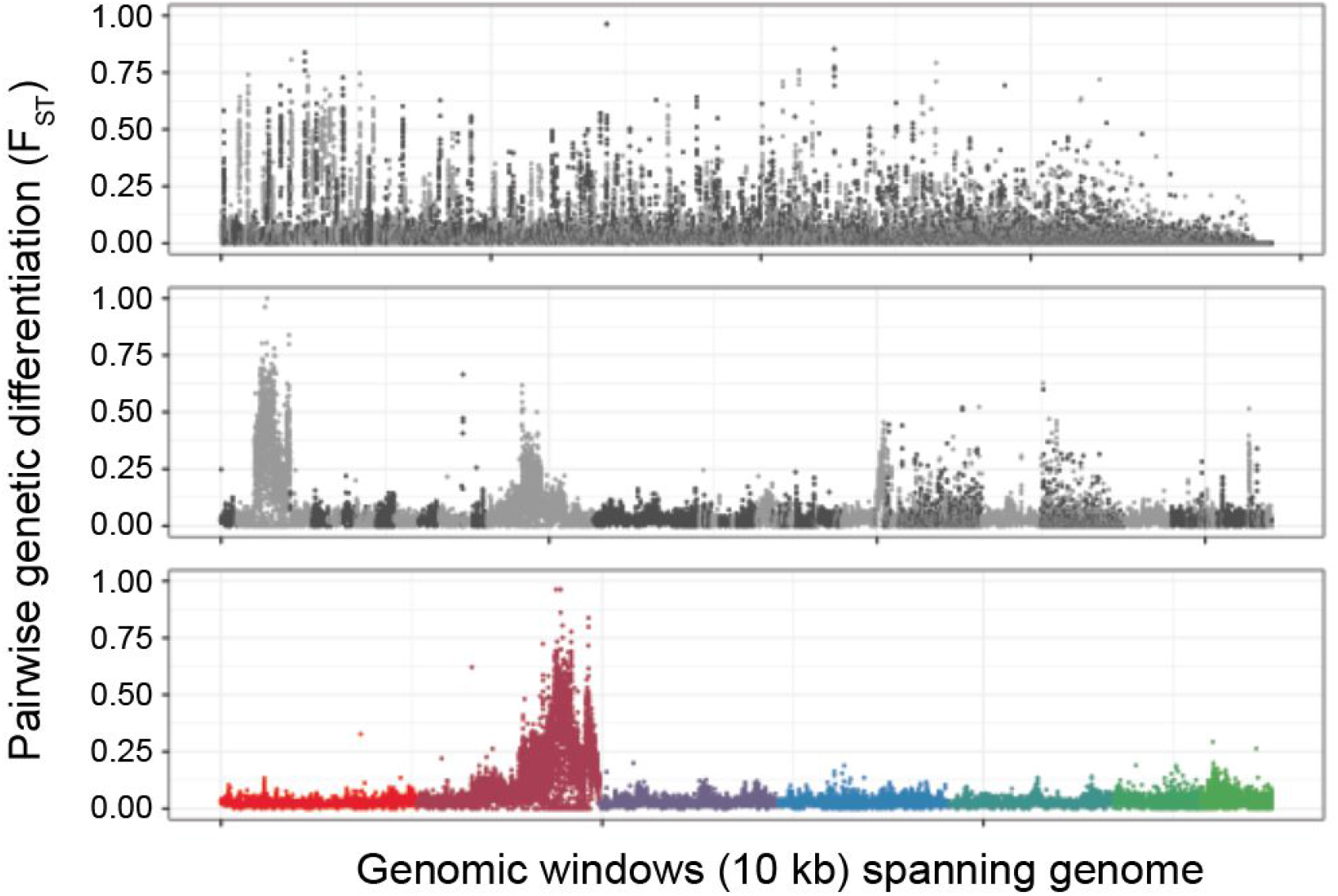
Impact of genome contiguity on the resolution of the introgression region. The same pairwise comparison – MHco3(ISE) *vs* MHco3/10.BC_4_.P_4_ – is presented using three versions of the *H. contortus* genome; the published V1 draft assembly [19] (top; N50 = 0.083 Mbp, N50(n) = 1,151, n = 23,860), an intermediate improved assembly (middle; N50 = 5.2 Mbp, N50(n) = 16, n = 6668), and the chromosomal-scale assembly presented here and elsewhere [21](bottom; N50 = 47.4 Mbp, N50(n) = 3, n = 8). The top and middle plots are coloured light and dark grey to reflect alternate contigs / scaffolds, whereas the bottom plot is coloured as in Fig 2A.

The extensive genome-wide diversity between parental populations re-emphasises that *H. contortus* is highly genetically diverse both within and between populations. It is clear, however, that short read mapping-based analyses exploiting a single reference genome will underestimate this diversity. In this study, we have used only a subset of the total variation present, i.e., only single nucleotide polymorphisms, to characterise the introgression; however, understanding the functional consequences of population-specific diversity will rely on a more comprehensive description of all of the variation that defines a population. As an alternative to a linear reference genome, the use of population genome graphs – a non-linear or branching reference that contains alternate paths representing known genetic variation – may be more suitable [51], and allow better characterisation of known variation that is too complex to be analysed using a linear reference [52]. This may be possible once a comprehensive analysis of population and perhaps global genetic diversity is made available [53]. Alternatively, *de novo* reference sequences from geographically diverse isolates may be required, ideally sequenced and assembled using long-read sequencing technology, to allow a more comprehensive description of genetic variation within a population while allowing large scale variation between reference populations to be characterised.

### Conclusion

Our genome-wide analysis of two genetic crosses between ivermectin resistant and ivermectin sensitive isolates has identified a major ivermectin resistance QTL shared between two genetically and geographically distinct populations of *Haemonchus contortus*. Traditional population genetic analyses highlighted the extensive genetic variation both between and within the parental populations used to construct the cross, potentially explaining why candidate genes identified using approaches that did not control for this genetic diversity have not been validated, and by using novel population genetic methods, we could better account for the crossing procedure and its impact on the outcome of the experiment. This work represents the most comprehensive analysis of the genetics of anthelmintic resistance in a parasitic nematode to date, and demonstrates the power of genetic crossing and a contiguous genome assembly to eliminate false positive genetic signals typically linked to resistance. Importantly, these data show that many of the previously proposed candidate genes are not involved in ivermectin resistance in these isolates, and should focus future efforts on identifying the causal variant within the QTL we identify.

## Methods

### Background of *H. contortus* populations and the original backcross

The MHco3(ISE), MHco4(WRS) and MHco10(CAVR) are *H. contortus* populations maintained and stored at the Moredun Institute. MHco3(ISE) was originally derived by multiple rounds of inbreeding of the SE population [46]. The precise provenance of SE is not clearly recorded but is thought to be originally obtained from East Africa. MHco10(CAVR) is derived from the Chiswick Avermectin Resistant (CAVR) strain, an ivermectin resistant field population originally isolated in Australia [54]. MHco4(WRS) is derived from the ivermectin resistant White River Strain (WRS) originally isolated from South Africa [14].

The experiment described here extends two previously-described backcross experiments. The construction, phenotypic validation, and initial microsatellite analysis of these backcrosses has been described previously [39], and is outlined in **Fig 1A and B.** Briefly, two independent crosses were performed in parallel: (i) MHco3(ISE) x MHco10(CAVR), and (ii) MHco3(ISE) x MHco4(WRS). In the first generation of each cross, female worms of the ivermectin resistant parental populations [MHco10(CAVR) or MHco4(WRS)] were crossed with male MHco3(ISE) worms, to generate lines designated MHco3/4 and MHco3/10, respectively (**Fig 1A**). After the first cross, the F_1_ generation females were mated to male MHco3(ISE), resulting in backcross generations designated MHco3/10.BC_n_ and MHco3/4.BC_n_, where *n* is the number of backcross generations (BC). This involved the recovery of immature worms from the abomasum at day 14 post infection by necropsy of donor sheep; worms were washed in physiological saline (0.85% NaCl) after which their sex was determined [55], followed by surgical transfer of 45-100 male MHco3(ISE) and 50-100 female F_1_ L_4_/immature adult worms into worm-free recipient lambs all within 2 h of the original collection. This process of backcrossing was repeated for a total of 4 generations, i.e., MHco3/10.BC_4_ and MHco3/4.BC_4_(**Fig 1B**). *In vivo* ivermectin selection was applied after mating and before collection of eggs to enrich for ivermectin resistant adults to be used in the subsequent round of the backcross. A controlled efficacy test was performed on the three parental-, MHco3/4.BC_4_, and MHco3/10.BC_4_ populations to determine the initial levels of ivermectin efficacy, and the degree of resistance acquired in the introgressed lines. After four rounds of backcrossing, a selection experiment was performed. MHco3/10.BC_4_ and MHco3/4.BC_4_ populations were used to infect recipient sheep, after which eggs were collected and in vitro culture to L_3_. These infective larvae were used to infect two sheep per population, one that received 0.1 mg/kg ivermectin and one that remained drug naive. At 7 days post treatment, treated and naive L_4_/immature adults were recovered by necropsy and stored for molecular analysis.

The backcrosses described above were extended by performing a further four rounds of *in vivo* ivermectin selection, during which mating within the BC_4_ population continued (**Fig 1C**). For each cross, progeny of the BC_4_ generation were cultured to L_3_ and used to infect a donor sheep, which was subsequently treated with ivermectin (0.1 mg/kg in round 1, followed by 0.2 mg/kg in following rounds). Eggs from ivermectin-treated survivor adults were collected post-treatment, cultured to L_3_, and used to infect a new donor sheep. L_4_/immature adults were collected by necropsy after rounds three (i.e., BC_4_.IVM.P_3_) and four (i.e., BC_4_.IVM.P_4_) of selection and stored for molecular analysis.

### Library preparation and sequencing

Whole genome sequencing was performed on the three parental populations (MHco3[ISE], MHco10[CAVR] & MHco4[WRS]), and from each cross, pre and post ivermectin treatment following four rounds of backcrossing (MHco3/10.BC_4_.noIVM, MHco3/4.BC_4_.noIVM, MHco3/10.BC_4_.IVM, MHco3/4.BC_4_.IVM), and after 3 and 4 rounds of additional selection after the backcross (MHco3/10.BC_4_.IVM.P_3_, MHco3/4.BC_4_.IVM.P_3_, MHco3/10.BC_4_.IVM.P_4_, MHco3/4.BC_4_.IVM.P_4_). Pools of male and female worms were included for the parental (n = 50-60 worms) and BC_4_ samples (n = 25-30 worms), whereas only female worms were used in the passage 3 & 4 samples (n = 60 worms). Genomic DNA from each pooled sample was prepared using a phenol chloroform extraction protocol as previously described. Sequencing libraries were prepared using a PCR-free protocol [56], and sequenced as described in **Table S1**. We generated approximately 6.14×10^11^ bp of sequence data, which equates to approximately 199.65× unmapped genome coverage per sample. Raw sequence data quality was analysed using FASTQC (*http://www.bioinformatics.babraham.ac.uk/projects/fastqc/*) and was assessed using MultiQC [57] prior to further processing.

### Mapping and variant analysis

Raw sequence data was mapped per lane to the version 3 reference genome (available here: ftp://ngs.sanger.ac.uk/production/pathogens/Haemonchus_contortus) using BWA-MEM [58](default parameters with -*k 15*). Samples for which multiple lanes of mapped data were available were merged, duplicate reads were marked and removed using *Picard v2.5.0* (https://github.com/broadinstitute/picard), and mapped perfect read pairs were extracted per sample (*samtools-1.3 view-f 2*).

Genome-wide variants were determined for each sample using samtools-1.3 mpileup (-*F 0.25-d 500*). Nucleotide diversity and Tajima’s D was determined using *npstat* [59], which required pileup files (which we derived from the mpileup generated above) that were split per sample per chromosome as input. We determined short-range linkage disequilibrium between pairs of variants in single and paired reads using *LDx* [60]. We compared these data with estimates of LD decay over genetic distance as proposed by Hill and Weir [60, 61]. *Popoolation2* [62] was used for the analysis of pairwise genetic diversity throughout the genome. Briefly, the mpileup file was converted into a synchronised file (*popoolation2 mpileup2sync.jar--min-qual 20*), after which indels + 5 bp (popoolation2 *identify-indel-regions.pl--min-count 2--indel-window 5*) and repetitive and difficult to map regions (separately identified by repeat masking the genome [http://www.repeatmasker.org/]) were excluded (*popoolation2 filter-sync-by-gtf.pl*). The synchronised file was used as input to determine pairwise F_ST_ which was calculated in 10 kbp windows throughout the genome (*popoolation2 fst-sliding.pl-pool-size 50 --window-size 10000 --step-size 5000 --min-count 4 --min-coverage 50 --max-coverage 2%*). Similarly, per base comparisons were determined using a Fisher’s exact test (*popoolation2 fisher-test.pl --min-count 4 --min-coverage 50 --max-coverage 2% --suppress-noninformative*).

Copy number variation between parental strains was determined using *cnv-seq* [63]. Best read mapping hits were determined per bam (*samtools-1.3 view-F 4 bam* / *perl-lane ‘print “ $F[2] \t$ F[3]”’ > bam.hits*), before read counts for pairwise comparisons determined in 10 kbp sliding windows using *cnv-seq.pl*. The R package *cnv* (v 0.2-8) was used to determine pairwise chromosome-wide normalised log_2_ ratios. Characterisation of structural variation in the parental populations was performed using the *speedseq sv* pipeline [64] to identify putative duplications, deletions and inversions in each population. This approach exploits all reads mapped using BWA-MEM, including split and discordant read pairs, which are subsequently scored using LUMPY and SVTyper. Conservative filtering was applied to retain only homozygous variants (1/1) with a minimum quality score of 100.

### Population genetic modelling

Bespoke single-locus and multi-locus models were used for population genetic inference from the data. Common to both methods, variants in the genome potentially associated with resistance were identified. At each locus in the genome, a nucleotide {A,C,G,T} was defined as existing in the susceptible parental population if it was observed at a frequency of at least 1% in that population. Loci in the resistant parental population were then identified for which exactly one nucleotide that did not exist in the susceptible population was observed at a frequency of at least 1%; without prejudice as to its phenotypic effect, this was denoted the ‘resistant’ allele at that locus, being associated with the resistant parental population.

#### Single-locus model

A single-locus population genetic model was then used to identify variants with frequencies that were inconsistent with selective neutrality. The neutral expectation was calculated using a Wright-Fisher evolutionary model to simulate the progress of a variant allele, modelling the probability distribution of the frequencies of homozygous resistant (*q*_1_), heterozygous (*q_h_*), and homozygous susceptible (*q*_0_) individuals throughout the course of the experiment under the assumption of selective neutrality. The Wright-Fisher model uses a simple multinomial process for propagation; if the next generation of the population contains *N* individuals, the probability of obtaining *n*_1_, *n_h_*, and *n*_0_ individuals with each diploid variant is given by:

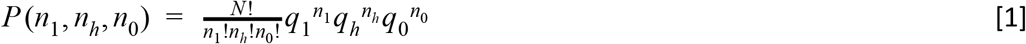

At the stages in the experiment where the population was comprised of eggs, we assumed *N* to be large, such that the processes of crossing and backcrossing could be described deterministically. Under these circumstances the genotype frequencies following a cross, denoted 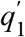, 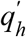, and 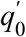, are given by:

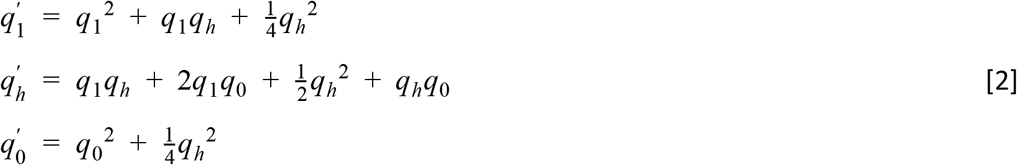

while the genotype frequencies following a backcross with the homozygous susceptible population are given by:

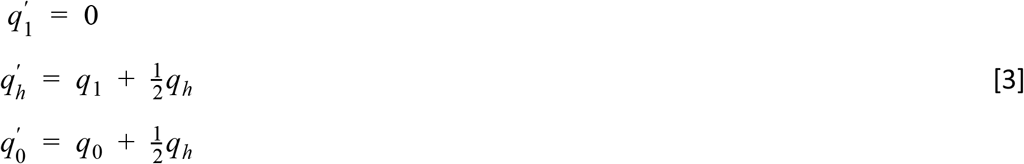

Evaluating this process gives a probability distribution for the frequency of the resistant allele at any given point in the experiment, dependent upon the number of resistant alleles in the initial resistant population. This number of resistant alleles, which we denote *n_r_*, is unknown; sequence data from the resistant parental population were used to generate a prior distribution for this value.

We first suppose that the frequency of the resistant allele in the resistant parental population is equal to some value, *p_r_*. Given that the initial experimental population contains 50 resistant worms, with 100 alleles, collected from this larger population, the distribution of *n_r_* is then given by:

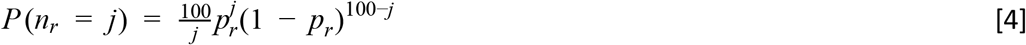

We now suppose that sequencing the resistant parental population gave *a_r_* resistant alleles at the locus in question out of a total read depth of *A_r_*. The initial frequency *p_r_* is unknown; however using the data an estimate can be made for this statistic, expressed as a distribution of the allele frequency. Specifically, under the assumption of a uniform prior, the underlying probability *p_r_* can be said to be distributed as a beta distribution with parameters *a_r_* and *A_r_ – a_r_* + 1. We therefore obtain the following distribution for the statistic *n_r_*:

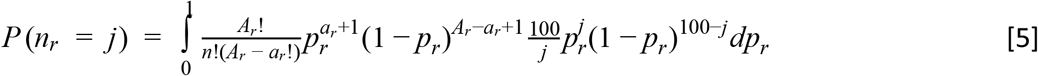

for any value of *j* between 0 and 100. Values of this distribution were calculated using numerical integration. This process generated the neutral expectation of the resistant allele frequency conditional on the observation of this frequency in the resistant population. If at the sampling point *t*, a total of *N_t_* worms were collected, the distribution of the number of worms *n_t_* with the resistant allele at that point is given by:

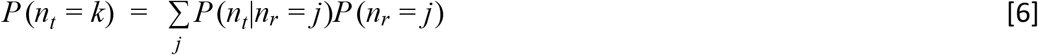

for all 0≤*k*≤*N_t_*

The extent to which observed allele frequencies were consistent with the neutral model was calculated using a likelihood model:

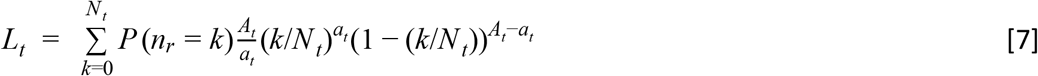

where at time *t, a_t_* resistant alleles from a read depth of *A_t_* were observed. A low likelihood *L_t_* indicates deviation from the neutral expectation; Bonferroni correction was used to identify significance at the 95% level. Data from each sample were analysed; four-fold non-neutral sites, being loci for which a significant likelihood was calculated from all four samples collected throughout each cross experiment, were identified.

#### Multi-locus model

A multi-locus model was developed to describe the manner in which allele frequencies would be expected to change over the course of the experiment; this model exploits information about the location of variant loci provided in the *H. contortus* reference genome, and the genome-wide map of recombination rate in the worms [21]; this map was inferred by characterising recombination breakpoints in F_1_ L_3_ progeny by whole genome sequencing, from which recombination rates were determined by comparing genetic distances between SNPs against their physical location in the genome. Where fixed genetic differences occur between the parental populations, the dynamics of adaptation in the resulting cross are relatively straightforward [48, 50]; this has been exploited to identify quantitative trait loci in yeast and malaria cross populations [49, 65]. Here a heuristic approach was used to identify fixed genetic differences between the parental populations before modelling evolution at these loci to identify the location of alleles conveying drug resistance.

Three filters were used to identify putatively fixed differences between parental populations. Firstly, we identified genomic sites for which the frequency of the resistant allele was 95% or greater in the resistant parent, requiring a read depth of at least 50x coverage for such sites to achieve a good level of statistical certainty. Secondly, we noted that, following the backcross performed at the start of the experiment, no locus can be homozygous for the resistant allele and as such, the resistant allele frequency can be no greater than 50% in the population; by way of reducing noise, we removed any locus with resistant alleles at a frequency of 60% or greater in the MHco3/10.BC_4_.noIVM sequence data. Thirdly, we noted that at homozygous separating sites in a population following a cross, the allele frequency will change smoothly over time; where there are *N* genomes in the population, the allele frequency, considered as a function of chromosome position, will change by 1/*N* wherever a crossover recombination event occurs within a genome. A diffusion model of allele frequency change, described in a previous publication [49], was used to identify allele frequencies across the genome that were consistent with this pattern. This analysis fits a posterior distribution to the allele frequencies across the genome, also inferring the extent of noise in the sequence data via a beta-binomial model intrinsic to the fitting process. Conservatively, sites no more than eight standard deviations from the mean of the posterior frequencies were included in the analysis. These approaches resulted in data from a total of 1,368 loci in the MHco10(CAVR) dataset, and from 219 loci in the MHco4(WRS) dataset, were retained; the full set of allele frequencies are shown in **S5 Fig**.

Considering these data, we used an individual-based Wright-Fisher simulation to model the outcome of the experiment, taking into account the backcrossing, selection, and bottlenecking according to the experimental design. This approach accounted for the stochastic nature of allele frequency changes due to the effect of repeated population bottlenecking. We suppose that selection acts in favour of the resistance allele at a given locus when worms are in a sheep being treated with ivermectin, such that the fitnesses of individuals that are homozygous for the resistant allele (*w*_1_), heterozygous for the resistant allele (*w_h_*), or homozygous for susceptible alleles (*w*_0_) are given by:

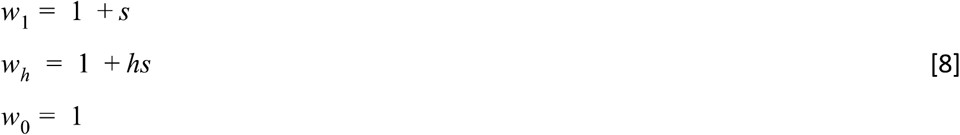

where *s* is the selection coefficient (fitness advantage of the homozygous resistant compared to homozygous susceptible) and *h* is the dominance coefficient (proportion of fitness change conveyed by a single copy of the allele). We assumed that all worms have equal fitness in the absence of the drug.

Our model is defined in terms of the manner in which selection acts upon the population, defined by the location in the genome of the selected allele, and the extent to which that allele conveys a fitness advantage to worms in the presence of drug treatment, this fitness advantage being characterised in terms of s and *h*. The likelihood of any given model was calculated using the beta-binomial likelihood function obtained from the diffusion process used above. Accounting for the stochasticity of the system, a set of at least 250 simulations were generated for each set of parameters, calculating the mean likelihood fit to the data across the simulations. A bootstrapping process was applied to quantify the uncertainty in this likelihood. Parameters giving the maximum likelihood fit to the data were identified.

Confidence intervals for the maximum likelihood locations of the selected allele were calculated. At the maximum likelihood position, we took the 250 simulations giving the maximum mean likelihood, calculating the mean of their likelihoods and the standard deviation of these values. We defined a threshold likelihood as the mean minus one standard deviation. For other allele positions, we also identified the parameters giving the maximum mean likelihood, constrained by allele position; we calculated a test likelihood equal to the mean plus one standard deviation of the replicate likelihoods generated by these parameters. Confidence intervals were then generated as the sites nearest the maximum likelihood position for which the test likelihood was lower than the threshold likelihood. We believe this gives a conservative estimate of the uncertainty in the location of the selected allele, given the stochasticity inherent in the experiment.

In an extension to this model, a two-driver scenario was considered, in which the resistant variant at each of two loci was under selection; to allow the exploration of model space in this case within reasonable computational time the assumption was made of additive selection at each locus. To distinguish this from the single-driver model, a requirement was imposed that the selected alleles be separated by at least 2 Mbp in the genome; this distance reflects the extent to which the process of recombination during the experiment allows the selective effects of distinct alleles to be observed.

## Declarations

### Ethics approval and consent to participate

All experimental procedures described in this manuscript were examined and approved by the Moredun Research Institute Experiments and Ethics Committee and were conducted under approved British Home Office licenses in accordance with the Animals (Scientific Procedures) Act of 1986. The Home Office licence numbers were PPL 60/03899 and 60/4421, and the experimental identifiers for these studies were E06/58, E06/75, E09/36 and E14/30.

### Consent for publication

Not applicable.

### Availability of data and material

The raw sequencing data generated during the current study are available in the European Nucleotide Archive repository (http://www.ebi.ac.uk/ena/) under the study accession number PRJEB2353 (**Table S1**). The reference genome assembly is available from ftp://ngs.sanger.ac.uk/production/pathogens/Haemonchus_contortus. Code used in this project is available from https://github.com/cjri/HCCross.

### Authors’ contributions

SRD performed the genomic analyses, and wrote the manuscript. CJRI performed the population modelling analyses, and wrote the manuscript. RL, DB, ER, AR, ED and AAM, performed the genetic crosses and collected and processed parasite material. AM, AT participated in genome curation. NH coordinated samples and sequencing. ED and MB provided supervision and guidance. NS, JAC and JSG participated in the experimental design and study supervision, and helped write the manuscript. All authors read and approved the final manuscript.

## Acknowledgements

We thank Pathogen Informatics and DNA Pipelines (WSI) for their support and expertise, and Guillaume Sallé and the Parasite Genomics team at WSI for constructive feedback on the manuscript. We are grateful to the Bioservices Division, Moredun Research Institute, for expert care and assistance with animals.

## Supplementary information captions

**S1 Table. Sample sequencing data archived at European Nucleotide Archive repository under the study accession PRJEB2353**

- See accompanying xlsx document

**S1 Fig.**
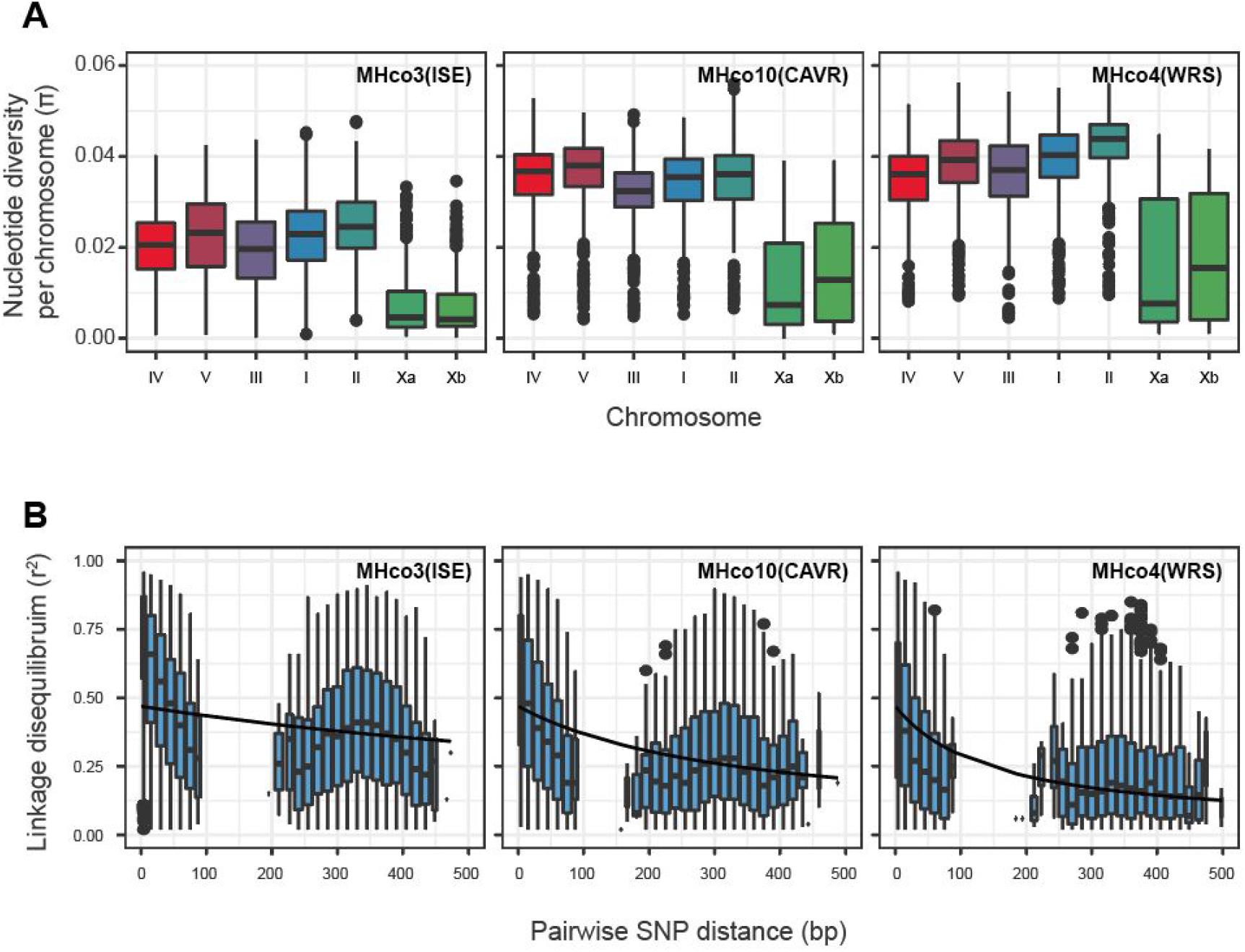
Characterisation of within-population diversity. **A.** Within population nucleotide diversity per chromosome, summarising genome-wide data presented in Fig 2C. Colours represent chromosomes as described in Fig 2A. **B.** Linkage disequilibrium between variants present in paired reads was estimated using LDx for each parental population. Line represents expected LD decay over genetic distance [60, 61].

**S2 Fig.**
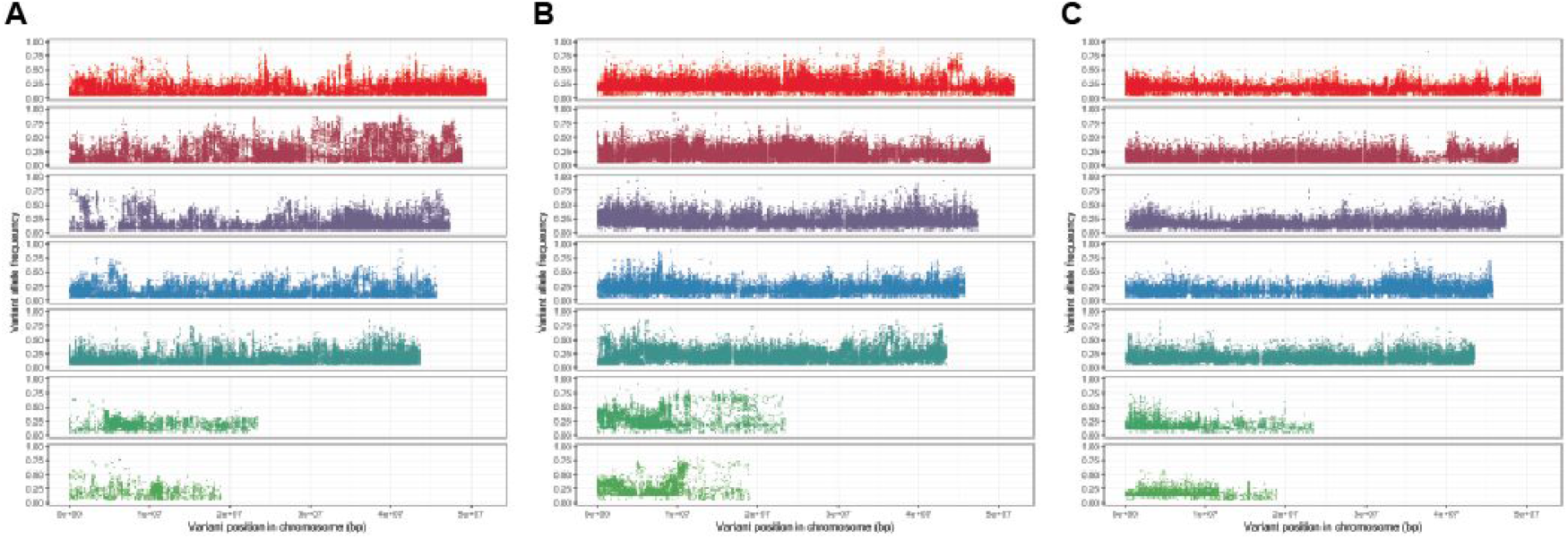
Distribution of “private” variant sites per parental population. **A.** MHco3(ISE). **B.** MHco10(CAVR). **C.** MHco4(WRS). Private sites were defined as having a frequency greater than 0.05 in the population of interest, but less than 0.05 in the two additional populations.

**S3 Fig.**
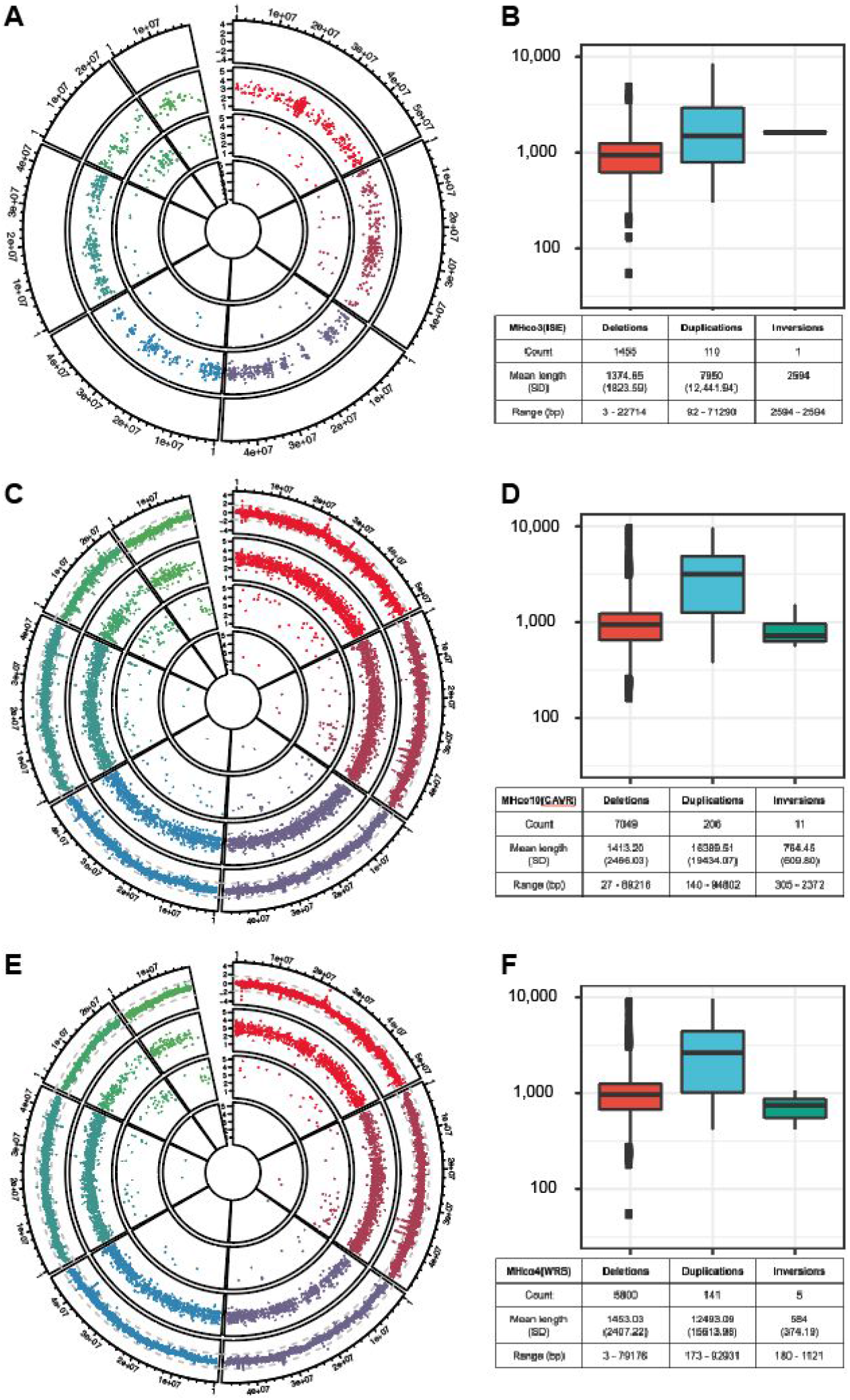
Copy number and structural variation in the parental lines. **A,B.** MHco3(ISE), **C,D.** MHco10(CAVR), **E,F.** MHco4(WRS). Data per circos plot **(A,C,E)** is orientated as follows; outer circle: CNV variation between MHco3(ISE) and each resistant parent. No CNV comparison was made in A; second circle: deletions; third circle: duplications; inner circle: inversions. The data presented in **A,C,E** is summarised in the boxplots (feature length distribution) and tables in **B,D,F.**

**S4 Fig.**
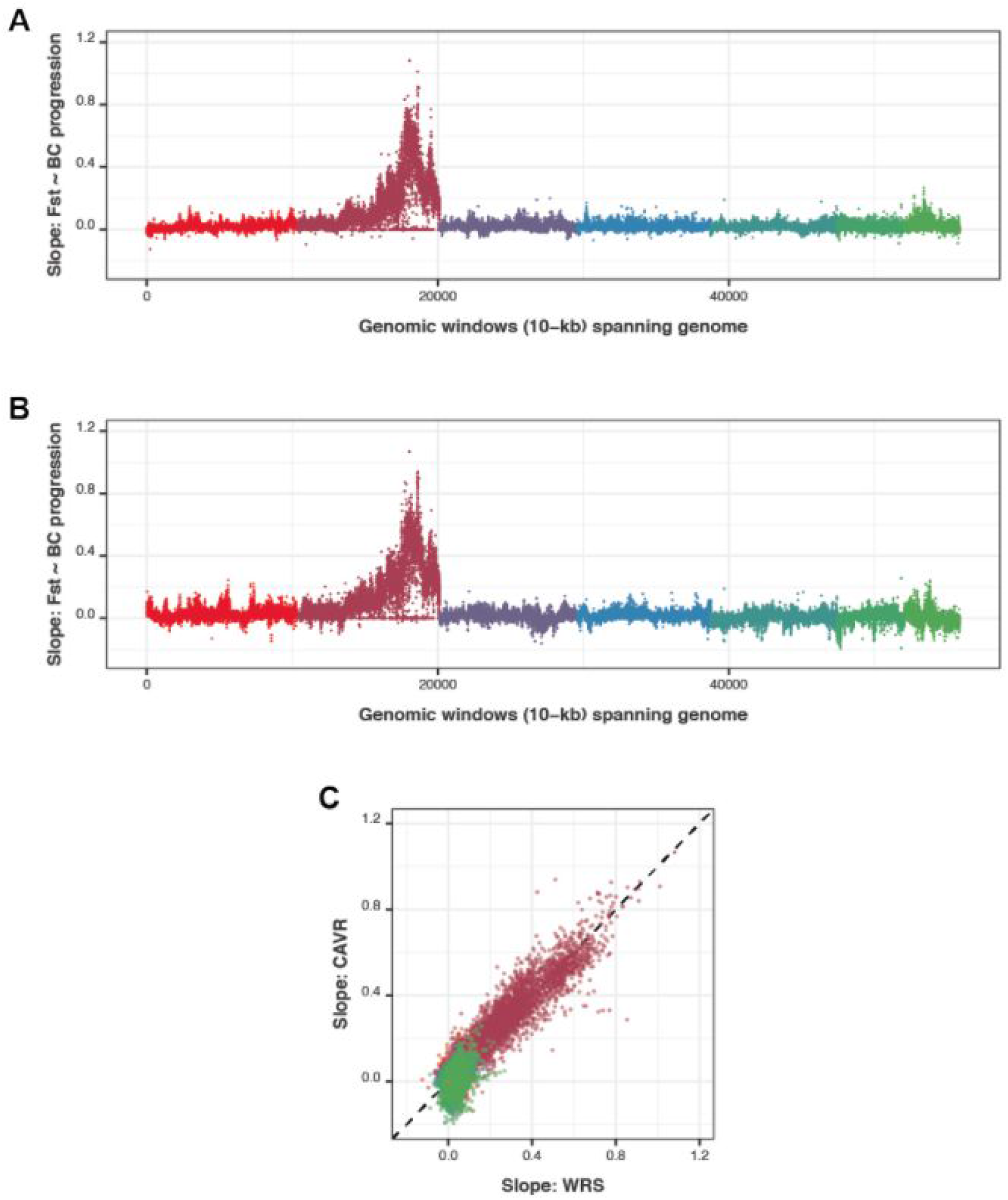
Summary of genome-wide change in F_ST_ throughout the backcross and subsequent passage. Linear regression between F_ST_ and the four sampling time points was performed for each 10 kbp window sampled across the genome for both MHco3/10 (**A**) and MHco3/4 (**B**). The slope of the regression was plotted. Panel **C** shows the correlation between the slopes (F_ST_ vs backcross progression) for MHco3/10 (A) and MHco3/4 (B). The dashed line represents x=y. Colours represent chromosomes as described in Fig 2A.

**S5 Fig.**
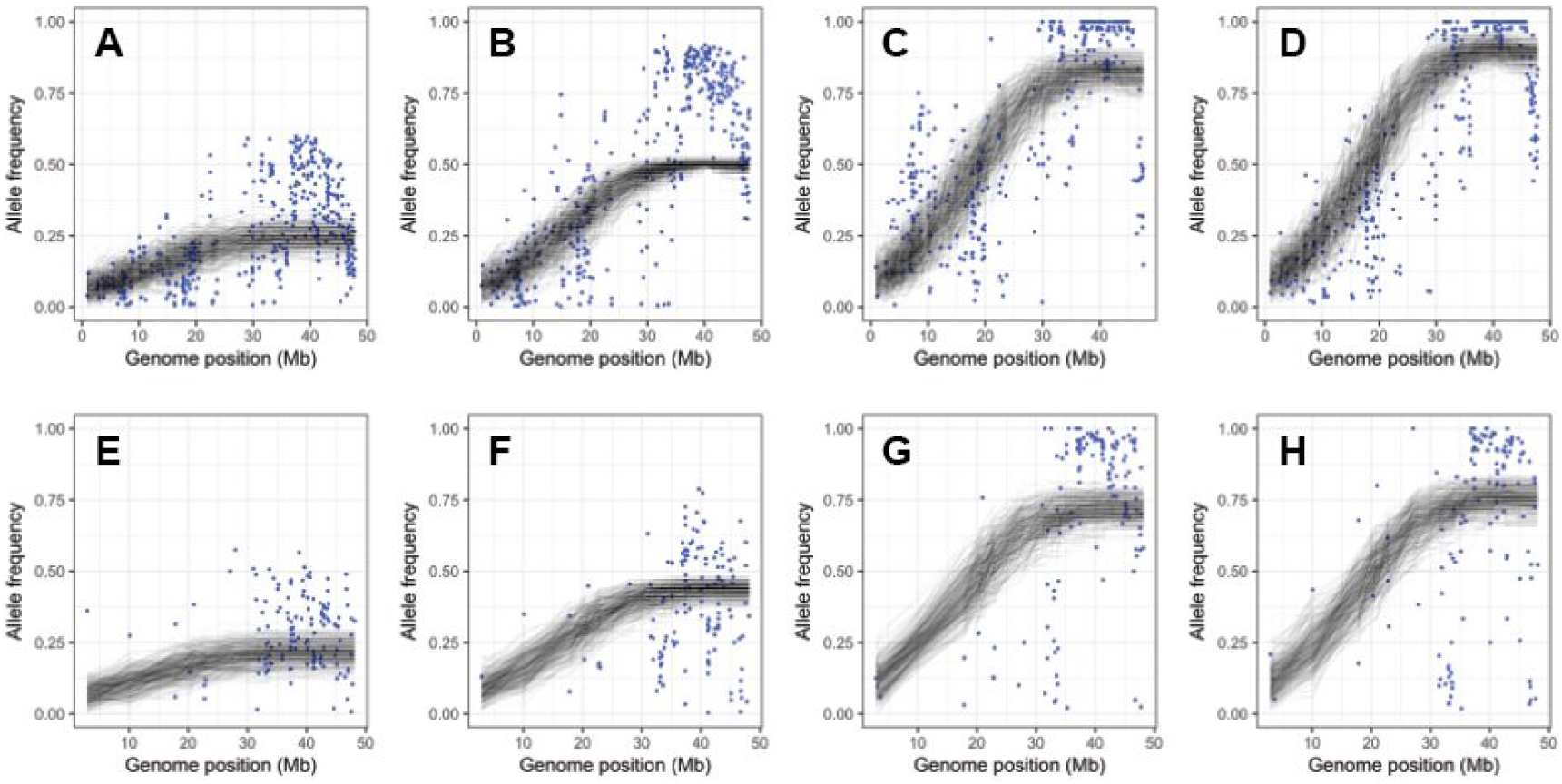
Fits between the model and the data for each data sample. Blue dots show filtered allele frequencies for putative segregating sites. The model fit is shown as gray lines; a distinct line is shown for each of the 250 replicate simulations run for the parameters generating the maximum likelihood fit. **A.** MHco3/10.BC_4_.noIVM. **B.** MHco3/10.BC_4_.IVM. **C.** MHco3/10.BC_4_.IVM.P_3_. **D.** MHco3/10.BC_4_.IVM.P_4_. **E.** MHco3/4.BC_4_.noIVM. **F.** MHco3/4.BC_4_.IVM. **G.** MHco3/4.BC_4_.IVM.P_3_. **H.** MHco3/4.BC_4_.IVM.P_4_.

**S6 Fig.**
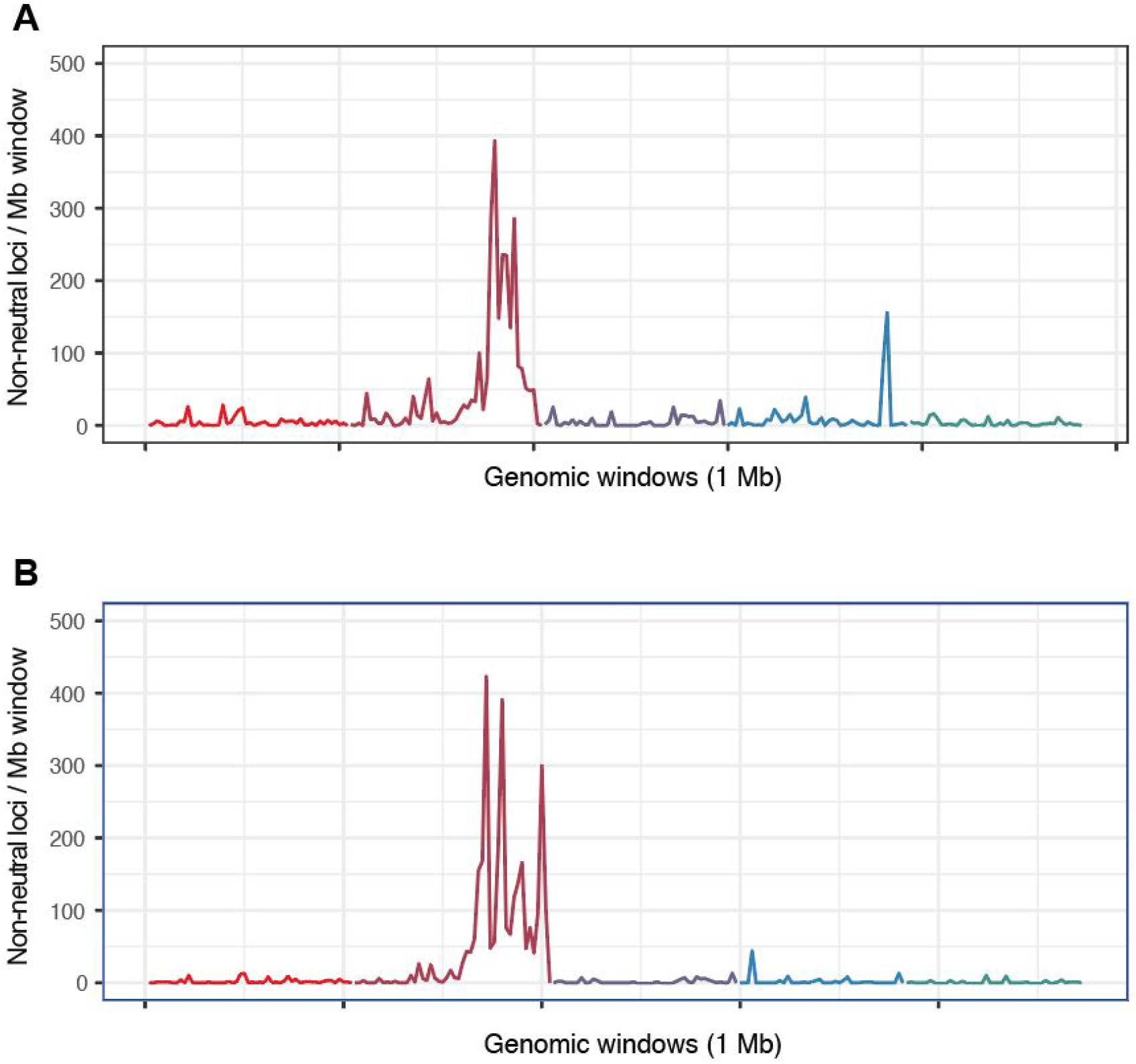
Location of significantly non-neutral loci identified using the single-locus population genetic model. Corresponding peaks in the location of significant sites can be seen in the MHco3/10 **(A)** and MHco3/4 datasets **(B).** A total of 70.6% of significant sites in the MHco3/10 dataset, and 90.6% of significant sites in the MHco3/4 dataset, were found in chromosome V. Data are binned in 1 Mbp windows spanning the genome. Colours represent chromosomes as described in Fig 2A.

**S7 Fig.**
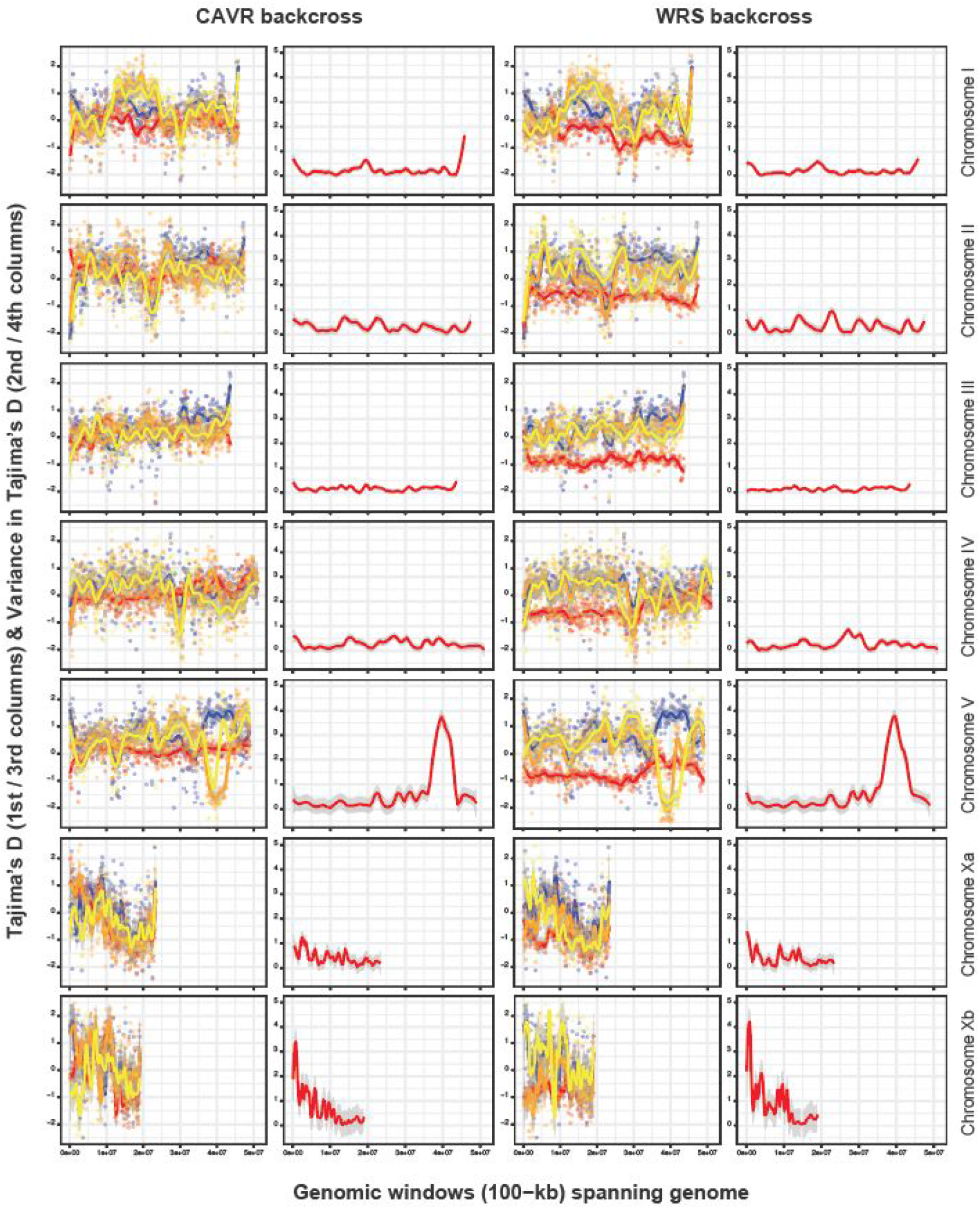
Analysis of Tajima’s D variation in each chromosome per cross. Comparison of Tajima’s D per chromosome between MHco3(ISE) parent (blue), MHco10(CAVR) (panel column 1; red) or MHco4(WRS) (panel column 3; red) and passages 3 (orange) and 4 (yellow) of the crosses. Tajima’s D was calculated using *npstat* in 100 kbp windows spanning the genome. The variance in the mean value of Tajima’s D among MHco3(ISE) and passages 3 and 4 – for which an increase in variance would suggest introgression and evidence of selection – was determined and is presented as smoothed line (red) in panel columns 2 and 4.

**S8 Fig.**
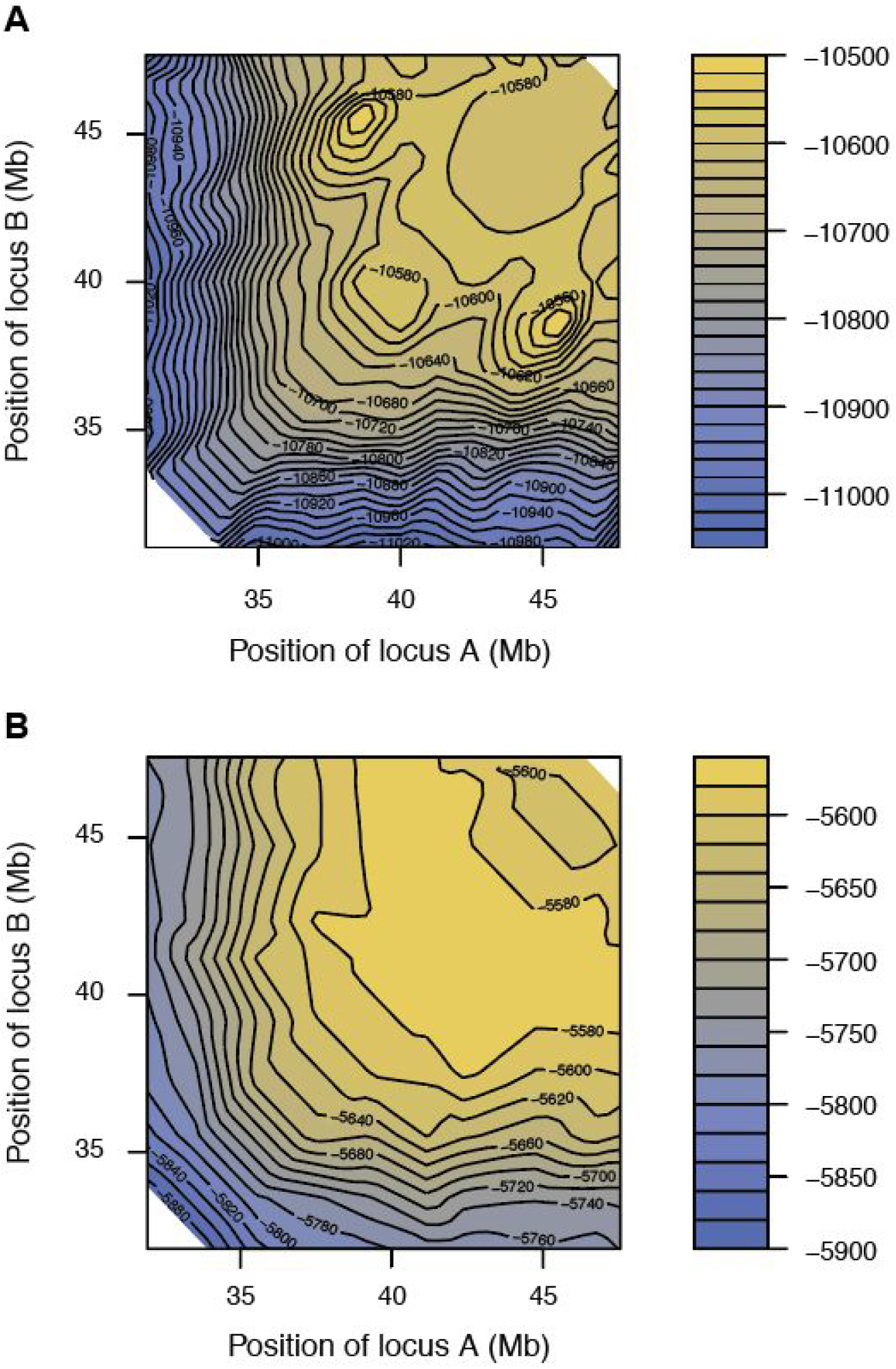
Contour maps of log likelihood scores derived from the two locus driver model. **A.** MHco10(CAVR). **B.** MHco4(WRS). The model was restricted to interactions between pairs of loci at least 2 Mbp apart.

**S9 Fig.**
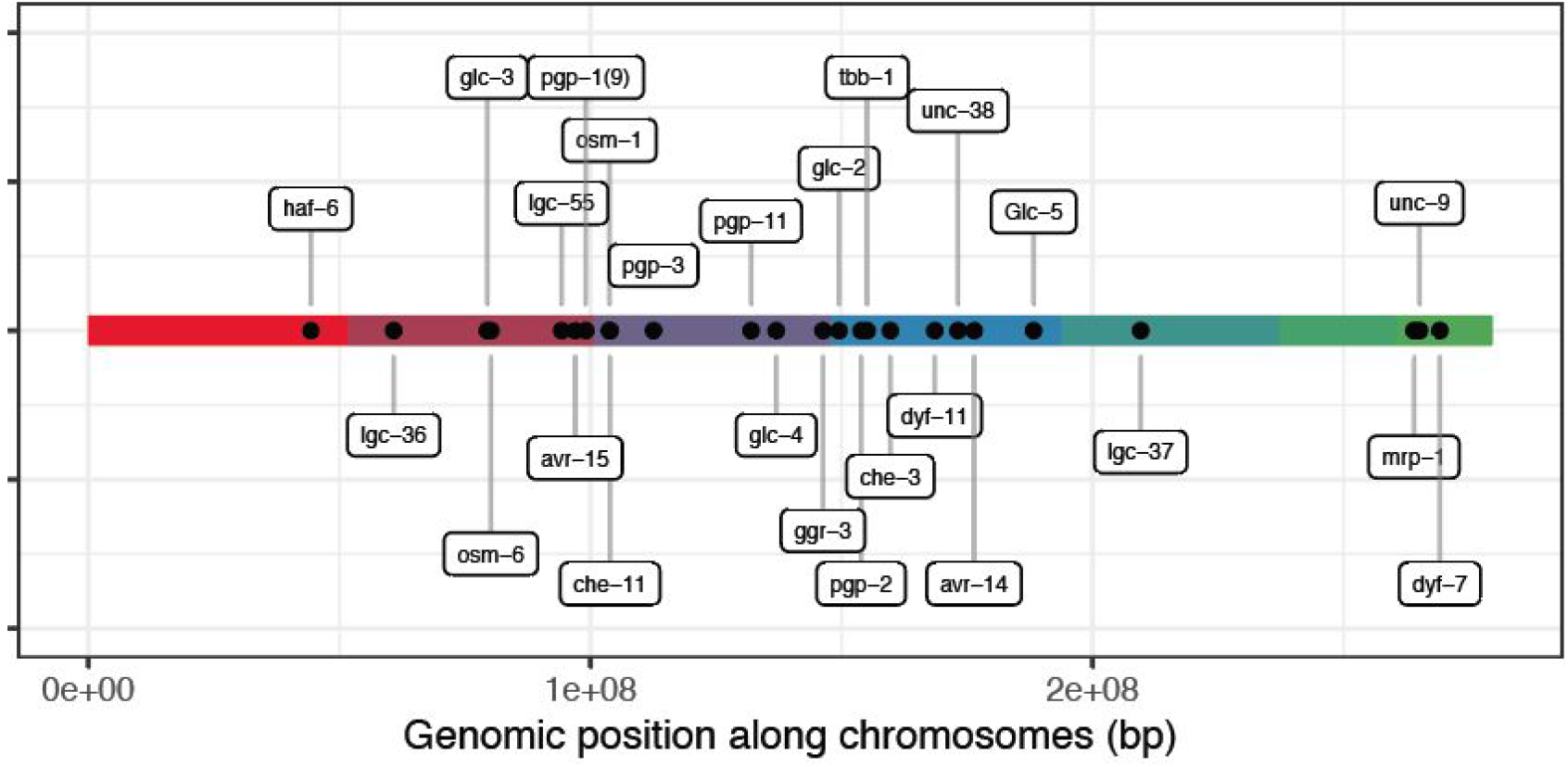
Relative position and location of candidate genes from the literature proposed to be associated with ivermectin resistance in *Haemonchus contortus* and / or *Caenorhabditis elegans*. Gene coordinates are presented in S2 Table. Colours represent chromosomes as described in Fig 2A.

**S2 Table.** Candidate genes from literature proposed to be associated with ivermectin resistance in *Haemonchus contortus* and / or *Caenorhabditis elegans*.

| Gene | GeneID <sup>3</sup> | Chromosome | Coordinates <sup>4</sup> | Reference |
| --- | --- | --- | --- | --- |
| *avr-14b* / *gbr-2* or *HcGluCl α3* <sup>2</sup> | HCOI00274000,<br>HCOI00274100 | I | 28352540 - 28362260 | (Williamson et al., 2011) |
| *avr-15* | HCOI00130400 | V | 45145104 - 45158307 | (Dent et al., 2000) |
| *tbb-1* | HCOI01967600 | I | 7026384 - 7029311 | (Eng et al., 2006; de Lourdes Mottier and Prichard, 2008) |
| *che-3* <sup>1</sup> | HCOI01710300 | I | 11666953 - 11695321 | (Dent et al., 2000) |
| *dyf-11* <sup>1</sup> | HCOI01514900 | I | 20494101 - 20505946 | (Dent et al., 2000) |
| *dyf-7* | HCOI01065900 | Xb | 8461163 - 8465904 | (Urdaneta-Marquez et al., 2014) |
| *ggr-3* | HCOI00098800 | II | 45626226 - 45639348 | (Rao et al., 2009) |
| *glc-1* <sup>1</sup> | NA |  |  | (Dent et al., 2000) |
| *glc-2* / *GluClb* <sup>2</sup> | HCOI00383000 | I | 1443439 - 1446121 |  |
| *glc-3* | HCOI00543300 | V | 27643249 - 27655188 | (Williamson et al., 2011) |
| *glc-4* | HPLM_0001206301 | II | 36314509 - 36331683 |  |
| *glc-5* / *HcGluClα* <sup>2</sup> | HCOI00617300 | I | 40196936 - 40206359 | (Blackhall et al., 1998) |
| *lgc-36* | HCOI02054500 | V | 9008092 - 9028181 |  |
| *lgc-37* / *HG-1* <sup>2</sup> | HCOI00977100 | III | 15741515 - 15762899 | (Blackhall et al., 2003) |
| *lgc-55* | HCOI00162900 | V | 42450788 - 42469177 |  |
| *osm-1*<sup>1</sup> | HCOI00970500 | II | 3105809 - 3126440 |  |
| *osm-3* | HCOI00164700,<br>HCOI00673700 |  |  |  |
| *osm-5*<sup>1</sup> | HCOI00498400,<br>HCOI00077900 |  |  |  |
| *pgp-1* / *pgp-9*<sup>2</sup> | HCOI00233200 | V | 47249715 - 47265234 | (James and Davey, 2009; Raza et al., 2016; van Wyk and Malan, 1988) |
| *pgp-2* / *pgp-a*<sup>2</sup> | HCOI00025600 | I | 5849505 - 5868032 | (Raza et al., 2016) |
| *pgp-11* | HCOI00622400 | II | 31321675 - 31344154 | (Raza et al., 2016) |
| *unc-38* | HCOI01939400 | I | 25118411 - 25125593 |  |
| *unc-7*<sup>1</sup> | HPLM 0000412001 |  |  | (Dent et al., 2000) |
| *unc-9*<sup>1</sup> | HCOI02075100 | Xb | 4410309 - 4424525 | (Dent et al., 2000) |
| *che-11* | HCOI00062600 | II | 3237676 - 3245201 |  |
| *osm-6* | HCOI00937900 | V | 28370676 - 28397433 |  |
| *pgp-3* | HCOI00117000 | II | 11888372 - 11913805 | (Raza et al., 2016) |
| *haf-6* | HCOI01007100 | IV | 44336050 - 44341372 | (Raza et al., 2016) |
| *pgp-13* / *pgp-12*<sup>1</sup> | HCOI01115500 | II | 10272077 - 10286519 | (David et al., 2018) |
| *mrp-1*<sup>1</sup> | HCOI01904000 | Xb | 3325452 - 3334565 | (James and Davey, 2009) |

1. *C. elegans* genes 2. Alternate names in the literature 3. Gene IDs correspond to the annotated gene in the published version of the MHco3(ISE) genome (Lainq et al., 2013). 4. Coordinates are approximate, as they represent the transfer of coordinates from the published version based on homology, and therefore may have changed depending on the conservation of the gene structure in the genome between the published and updated genome version used here.

**S10 Fig.**
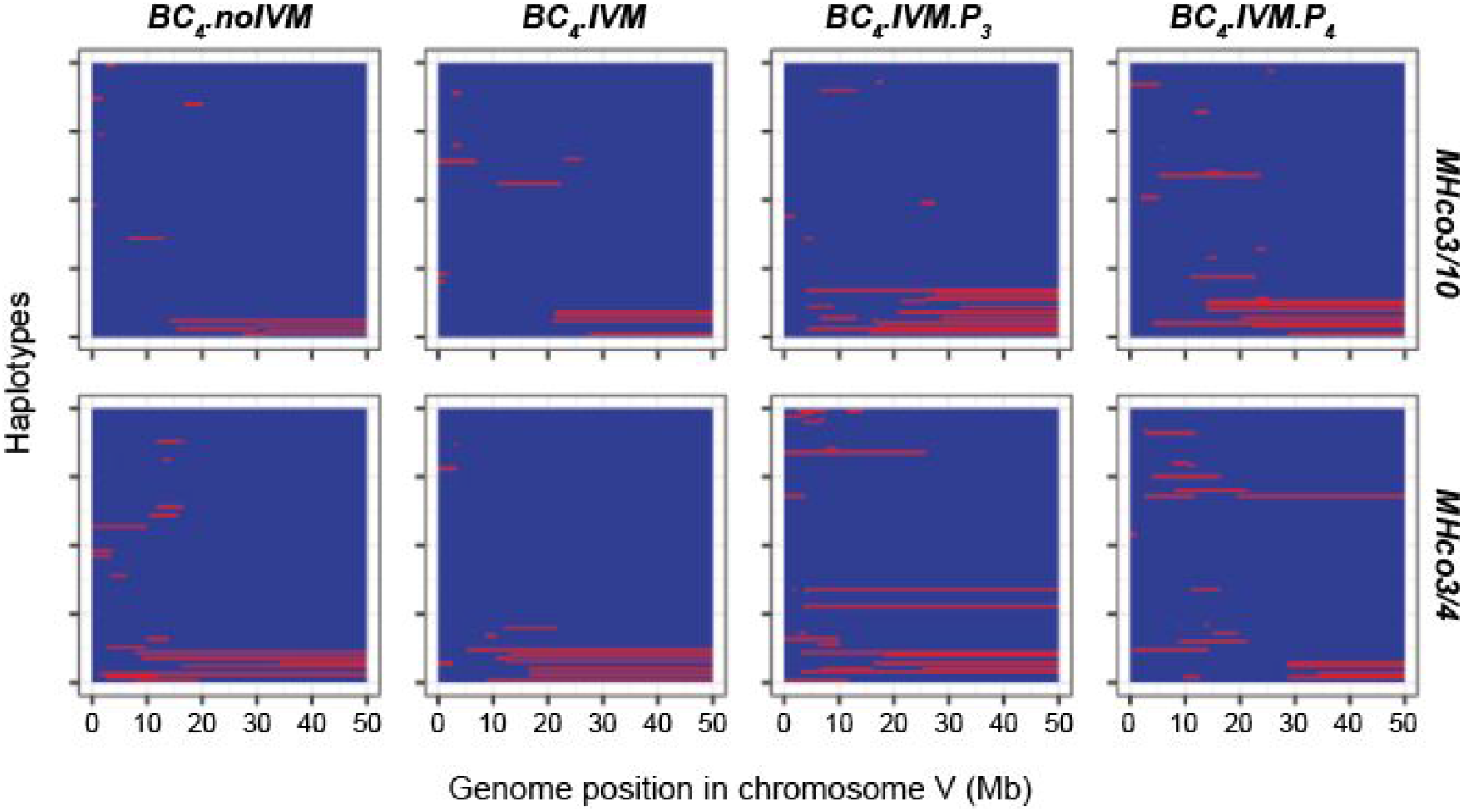
Haplotype structure of chromosome V in an example output from the model under neutral evolution. Segments of genome from the resistant parent are shown in red, while segments of genome from the susceptible parent are shown in blue. The repeated backcross removes most of the resistant genotypes from the population.

